# The transcriptional repressor Zfp125 modifies hepatic energy metabolism in response to fasting and insulin resistance

**DOI:** 10.1101/2020.07.02.185165

**Authors:** Gustavo W. Fernandes, Barbara M. L. C. Bocco, Tatiana L. Fonseca, Federico Salas-Lucia, Olivia Nickel, Samuel C. Russo, Balázs Gereben, Isis C. Kettelhut, Antonio C. Bianco

**Affiliations:** Section of Endocrinology, Diabetes and Metabolism, Department of Medicine, University of Chicago, Chicago IL; Laboratory of Molecular Cell Metabolism, Department of Endocrine Neurobiology, Institute of Experimental Medicine, Budapest, Hungary; Department of Biochemistry and Immunology, Ribeirão Preto Medical School, University of São Paulo, Ribeirão Preto, SP, Brazil

**Author notes:** authors contributed equally to this work (Gustavo W. Fernandes, Barbara M. L. C. Bocco). Disclosures: AB is a consultant for Allergan Inc, Synthonics Inc and BLA Technology LLC; the other authors have no disclosures. Corresponding author: Antonio C. Bianco, MD, PhD, Section of Adult and Pediatric Endocrinology, Diabetes and Metabolism 5841 S. Maryland Ave., Chicago, IL, 60637 USA.

**Keywords:** Zfp125, ketogenesis, fatty acid oxidation, gluconeogenesis, glucagon, insulin resistance

## Abstract

Zfp125 is a transcriptional repressor that inhibits hepatic VLDL secretion. Here we show that mice with liver-specific Zfp125 knockdown exhibited lower respiratory quotient, reduced glycemia and pyruvate-stimulated liver glucose output, and higher levels of β-hydroxyl-butyrate. Microarray and ChIP-seq studies identified Zfp125 peaks in the promoter of 135 metabolically relevant genes, including genes involved in fatty acid oxidation and ketogenesis, e.g. Ppara, Cpt1a, Bdh1 and Hmgcs2. Repression by Zfp125 involved recruitment of the corepressors Kap1 and the histone methyl transferase Setdb1, increasing the levels of H3K9me3, a heterochromatin marker of gene silencing. The resulting increase in acetyl-CoA levels accelerated gluconeogenesis through allosteric activation of pyruvate carboxylase. Zfp125 knockdown in isolated mouse hepatocytes amplified the induction of ketogenesis by glucagon or insulin resistance, whereas the expression of key gluconeogenic genes Pck1 and G6pc was amplified by Zfp125. These findings place Zfp125 at the center of fuel dysregulation of type 2 diabetes.

## Introduction

The liver is central in controlling the availability of energy substrates that support homeostasis. Dysregulation of these processes can lead to serious consequences such as excessive triglycerides (TG) accumulation, considered the major hallmark of nonalcoholic fatty liver disease (NAFLD) (Fon Tacer and Rozman, 2011). NAFLD affects more than 100 million individuals in the United States, and is commonly associated with obesity and insulin resistance (IR) (Browning and Horton, 2004; Fon Tacer and Rozman, 2011). The latter is key to increase TG deposit due to accelerated *de novo* lipogenesis (DNL) (Donnelly et al., 2005; Fon Tacer and Rozman, 2011; Lambert et al., 2014) and uptake of non-esterified fatty acids (NEFAs) (Sunny et al., 2011).

Caloric/carbohydrate restriction or fasting also accelerate liver uptake of NEFA, but in this case the faster rates of β-oxidation generate acetyl-CoA that is diverted from the TCA cycle and used in the ketogenesis pathway (Browning et al., 2012; Browning et al., 2008). Accelerated ketogenesis results in the hepatic release of β-hydroxybutyrate (BHB) (Balasse, 1979), the main ketone body that is utilized as energy source by peripheral tissues, such as the skeletal muscle (Fletcher et al., 2019). The opposite is observed in NAFLD, with acetyl-CoA primarily being used in the TCA cycle, despite increased oxidative metabolism. In fact, as the hepatic TG content increases, there is a slowdown in the production of ketone bodies in human liver (Fletcher et al., 2019). In NAFLD, reduced hepatic output of BHB is associated with accelerated acetyl-CoA oxidation in the TCA cycle and elevated rates of gluconeogenesis, illustrating how the liver increases oxidative metabolism instead of disposing of the excess of NEFAs as ketone bodies (Fletcher et al., 2019).

Zfp125 is a liver Kruppel associated box (KRAB) containing zinc finger protein (ZFP) that functions as a transcriptional repressor, slowing down lipid transport and hepatic VLDL secretion, leading to liver steatosis (Fernandes et al., 2018a; Fonseca et al., 2019; Fonseca et al., 2015). Here we show that Zfp125 inhibits fasting-induced ketogenesis by targeting and repressing critical genes involved in fatty acid oxidation and synthesis of BHB. Food availability and insulin secretion normally terminate this effect by inhibiting Zfp125 expression. However, Zfp125 expression remains elevated in states of IR, limiting activation of ketogenesis and contributing to the higher rates of gluconeogenesis typically seen when insulin signaling is impaired.

## Results

### A mouse with liver-specific Zfp125 knockdown exhibits increased contribution of fatty acid oxidation to energy expenditure

To explore the physiological role of Zfp125, we admitted mice to a Comprehensive Lab Animal Monitoring System (CLAMS). After a period of acclimatization, liver-specific Zfp125 knockdown was achieved through i.p. injections of liposomes containing Zfp125 siRNA during 5 days; control animals received liposomes loaded with scrambled RNA (Fig. 1A; Fig. S1A) (Fernandes et al., 2018b). Zfp125 knockdown decreased hepatic Zfp125 mRNA levels by ~45% (Fig. 1B) and markedly reduced Zfp125 protein levels (Fig. S1B-C), without affecting Zfp125 mRNA in epididymal white adipose tissue (eWAT) and soleus muscle (Fig. S1 F-G). No differences were observed in food intake, body weight or liver weight, eWAT or lean carcass (skeletal muscle and bones; Fig. S1 H-L) weights. Even though the circulating levels of NEFA and insulin were not affected (Table 1), Zfp125 siRNA reduced the respiratory quotient (RQ) *vs* control animals (AUC p<0.05; Fig. 1A). These animals also exhibited a ~2-fold elevation in plasma BHB (Fig. 1C) and a ~15% reduction in plasma glucose levels (Fig. 1D), while hepatic glycogen content remained unaffected (Fig. S2B-C).

**Figure 1.**
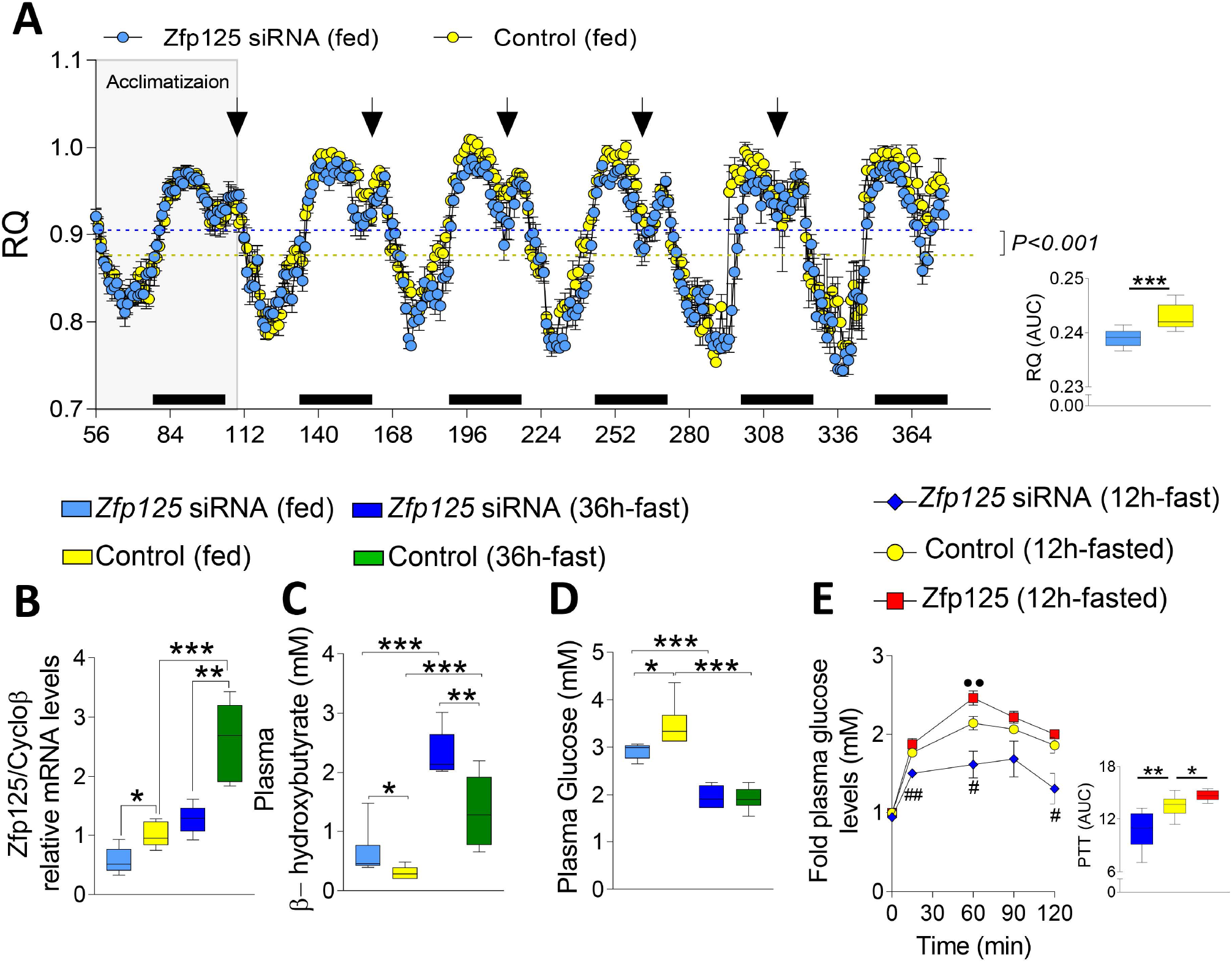
Effects of liver-specific Zfp125 knockdown on metabolic parameters. **(A)** Fluctuation of respiratory quotient (RQ) of adult mice while admitted to a CLAMS; after acclimatization, mice received i.p. injections of liposome suspension containing Zfp125 siRNA at 0h, 24h, 48h, 72h and 96h (arrows); control animals received liposomes loaded with scrambled RNA; the area under curve (AUC) is shown in the far right; some animals were fasted for 36h immediately before killing (Fig. S2A); at the end of the experiment, all mice were killed and hepatic Zfp125 mRNA levels as assessed by RT-qPCR are shown in **(B)**; plasma BHB levels are shown in **(C);** plasma glucose levels are shown in **(D)**; **(E)** pyruvate tolerance test (PTT) performed in mice treated as in **(A)** and fasted for 12h; plasma glucose levels were assessed over a period of 2h; AUC is shown on the far right; values are the mean ± SEM of 6-12 independent samples; *P<0.05, **P<0.01, ***P<0.001 as indicated; •• P<0.01 vs Control (12h-fasted); #P<0.05 vs Control (12h-fasted). Differences calculated by Student’s t-test in A, and One-way ANOVA was used in B-E.

**Table 1:** Different parameters in the plasma of fed/fasted mice with liver-specific Zfp125 siRNA.

| *Plasma parameter* | *Non-target siRNA (fed)* | *Zfp125 siRNA (fed)* | *Non-target siRNA (36h-fasted)* | *Zfp125 siRNA (36h-fasted)* |
| --- | --- | --- | --- | --- |
| NEFA (mM) | 0.54 ± 0.3 | 0.83 ± 0.2 | 1.4 ± 0.5•• | 1.5 ± 0.1 ** |
| Acetoacetate (mM) | 0.04± 0.01 | 0.04 ± 0.01 | 0.06 ± 0.02 | 0.05 ± 0.02 |
| Insulin (ng/ml) | 0.48± 0.01 | 0.51± 0.06 | 0.13± 0.04••• | 0.11± 0.01*** |

Values in this table are shown as mean ± SD of 4 independent samples. ••P<0.01, •••P<0.001 vs non-target siRNA (fed); \*\*P<0.01, \*\*\*P<0.001 vs Zfp125 siRNA (fed); differences calculated by one-way ANOVA followed by Student-Newman-Keul's test.

To test whether Zfp125 plays a role in hepatic energy output, mice with liver-specific Zfp125 knockdown were fasted for 36h, which prevented the usual increase in Zfp125 mRNA levels caused by fasting (Fig. 1B); Zfp125 protein levels followed a similar pattern (Fig S1D-E). Fasting prompted a marked drop in RQ values in both control and Zfp125 siRNA mice, indicating marked acceleration of fatty acid oxidation (Fig. S2A). However, the induction of plasma BHB levels by fasting was ~70% higher in the animals with liver-specific Zfp125 knockdown (Fig. 1C), without affecting acetoacetate levels (Table 1). This occurred even as the elevation in NEFA levels (1.9-2.5-fold) was similar in both groups (Table 1). Fasting reduced hepatic glycogen content (Fig. S2D-E) and plasma glucose levels by approximately 30-45% (Fig. 1D) and plasma insulin by ~80% in both groups (Table 1). Notably, fasted mice that received Zfp125 siRNA lost ~50% less body weight than those in the control group (Fig. S2F) possibly as a result of the borderline higher lean carcass weight (p=0.06; Fig. S2I); no differences were observed in liver weight and eWAT between groups (Fig. S2G-H).

To test whether Zfp125 affected gluconeogenesis we performed a pyruvate tolerance test for 2h in mice that were fasted for 12h. Hepatic glucose production (HGP), as estimated by plasma glucose levels after administration of pyruvate, was ~20% lower in mice with liver-specific Zfp125 knockdown (AUC, P<0.01; Fig. 1E). The opposite was seen in mice with liver-specific overexpression of Zfp125, that exhibited a ~10% higher glucose levels after pyruvate administration (Fig. 1E).

### Zfp125 knockdown accelerates ketogenesis and slows down gluconeogenesis in AML12 liver cells

We next developed a cell model using the AML12 liver cell line to study the role of Zfp125 in ketogenesis and gluconeogenesis during conditions of transient knockdown (Zfp125 siRNA) or stable expression of Zfp125; the latter mimics the upregulation in Zfp125 seen during fasting, approximately 2.5-fold (Fig. 2A). To study ketogenesis we took advantage of the fact that hepatocytes cannot take up or metabolize ketone bodies (Laffel, 1999). Thus, BHB level in the medium was measured over time when only nonessential amino-acids (NEAA) were present or when NEAA were supplemented with 2mM sodium octanoate. In the presence of NEAA, the level of BHB in the medium increased over time, reaching ~300uM after 24h and ~350uM by 48h (Fig. 2B). In cells treated with Zfp125 siRNA, the accumulation of BHB reached ~470uM in the first 24h and ~580uM by 48h (Fig. 2B). At the same time, cells stably expressing Zfp125 exhibited BHB levels of ~100uM and ~190uM after 24 and 48h, respectively (Fig. 2B). Similar results were obtained when sodium octanoate was added to the medium (Fig. 2C). The total accumulation of BHB reached ~590uM in cells treated with Zfp125 siRNA and only ~190uM in cells stably expressing Zfp125 (Fig. 2C).

**Figure 2.**
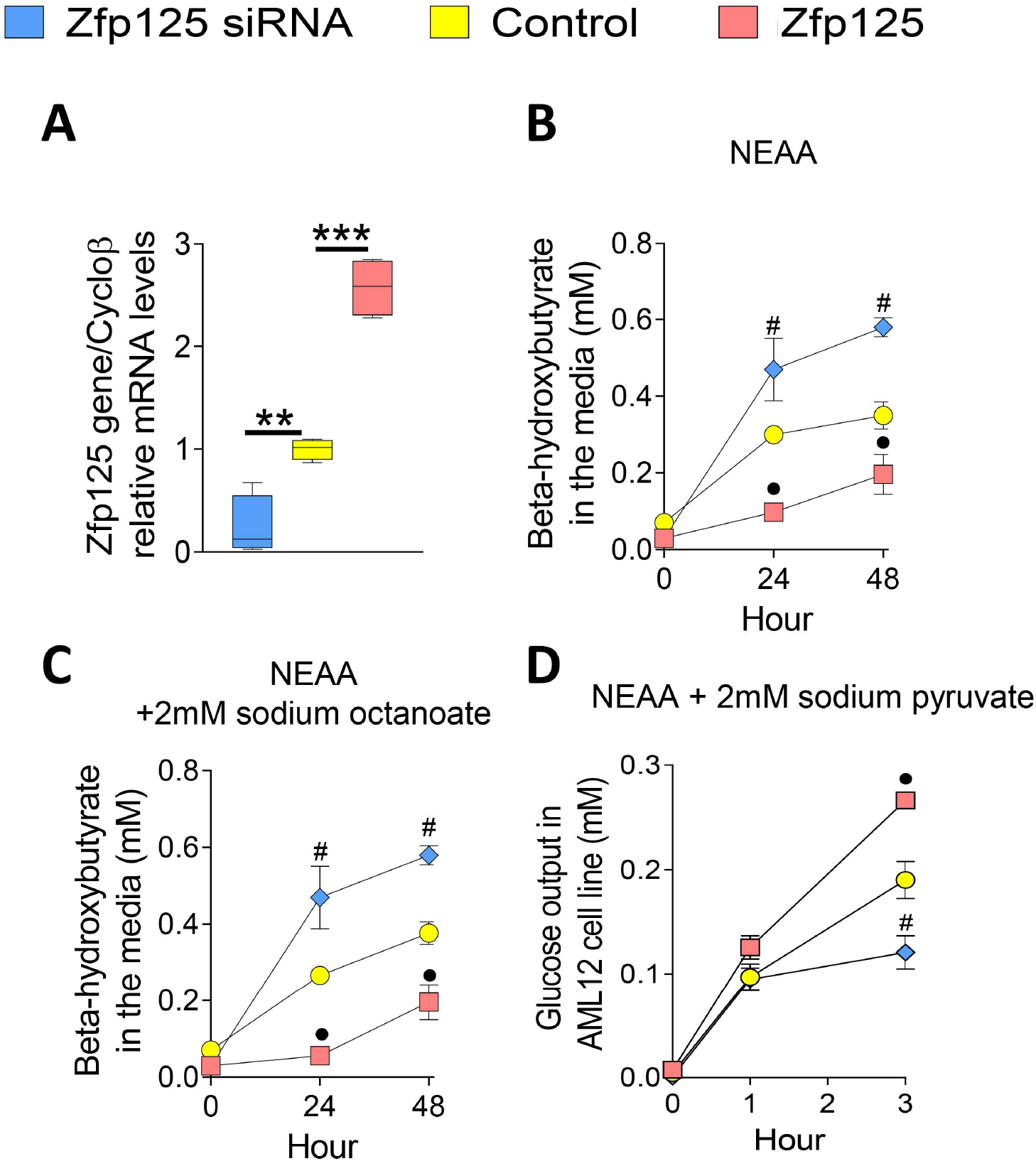
Effects of Zfp125 on ketogenesis and gluconeogenesis in ALM12 cells. **(A)** Zfp125 mRNA levels in AML12 cells stably expressing Zfp125, transiently transfected with Zfp125 siRNA or scrambled RNA; (**B**) BHB levels in the medium of cells incubated with NEAA; (**C**) same as (**B**) except that medium contained NEAA and 2mM sodium octanoate; (**D**) same as (**B**) except that cells were incubated with glucose-free medium supplemented with 2mM pyruvate; glucose measurements were obtained at 0h, 1 h and 3h; values are the mean ± SD of 3-1 2 independent samples. **P < 0.01, *** P<0.001 as indicated; • P<0.05 vs control; # P<0.05 vs control; differences calculated by one-way ANOVA followed by Student-Newman-Keul’s test in A-D; NEAA is nonessential amino-acids.

Next, we used the AML12 cell model to test whether glucose output was affected by Zfp125. Cells stably expressing Zfp125 and cells transfected with Zfp125 siRNA were pre-incubated in glucose-free medium during 3h for glycogen depletion, and transferred to glucose-free medium supplemented with 2mM pyruvate. In control cells, glucose levels in the medium increased over time, reaching ~90uM after 1h and ~190uM by 3h (Fig. 2D). Zfp125 knockdown limited the elevation in glucose levels to 120uM by 3h whereas in the cells stably expressing Zfp125 glucose levels reached ~260uM (Fig. 2D).

### Zfp125 is a transcriptional repressor of metabolically related genes

Through *in silico* analyses and PCR we identified Zfp125 as a 38-kDa protein containing 10 C2H2 zinc fingers connected to a KRAB box domain, the essential features of KRAB zinc finger proteins (Fig. S3A)(Sripathy et al., 2006); Zfp125 is highly homologous to its human counterpart ZNF670 [Fig. S3A-B; S4A-B; (Fernandes et al., 2018a)]. Using α-Zfp125 we identified Zfp125 in the cytoplasm and in the cell nucleus of mouse hepatocytes (Fig. S5A). The cytoplasmic staining was more intense in periportal hepatocytes, and not present in endothelial, ductal or arteriolar cells (Fig. S5A). These features were also seen in primary mouse hepatocytes in culture (Fig. S5B). Nuclear Zfp125 was also detected in western blots of AML12 cells stably expressing Zfp125 (Fig. S5C), and after immunoprecipitation (Fig. S5D).

We next performed chromatin immunoprecipitation followed by deep sequencing (ChIP-seq) in AML12 stably expressing Zfp125 to obtain an unbiased assessment of chromatin areas directly targeted by this protein. Three independent samples of Zfp125-AML12 ChIP were used to generate aligned reads of ~3-4 GB. Genome wide distribution of the ChIP-seq identified 644,831 peaks (18,294 genes; Fig. 3A and Table S1), with about half located in intronic regions, ~34% located in intergenic regions, ~13% in exonic regions and ~1.5% in promoter regions (Fig. 3A). We focused on the 9,944 Zfp125 ChIP peaks located in gene promoters, where their density increased with proximity to the transcription start sites (TSS; 6,724 genes; Table S2) (Fig. 3B). Two novel putative consensus-binding motifs for Zfp125 were identified (Fig. 3C-D). A heatmap of Zfp125 binding motifs *vs* Zfp125 ChIP peaks shows 80% overlap (Fig. 3E), which is in keeping with our working model that Zfp125 is a transcriptional factor. That the scope of action of Zfp125 includes regulating metabolically related genes was revealed through pathway analysis (Partek-Flow) of the genes containing Zfp125 ChIP peaks in the promoter (Table S3-4).

**Figure 3.**
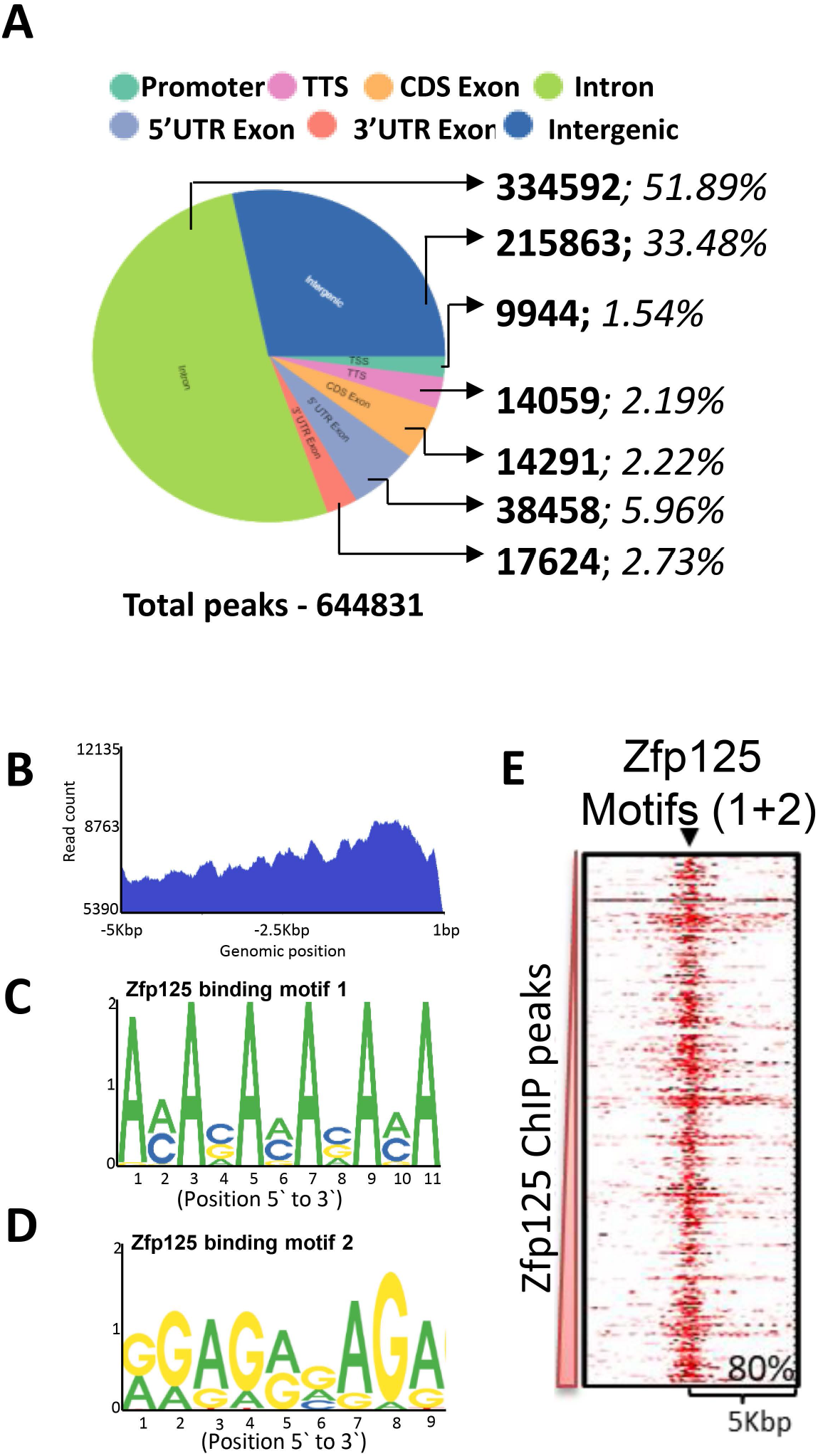
ChIP seq using αZfp125 in AML12 cells. **(A)** Distribution of Zfp125 ChIP peaks after GENECODE GENE (release M24) annotation; the breakdown of the different locations is indicated and include promoter region, transcriptional terminal site (TTS), coding sequence exons region (CDS Exon), introns, 5’ UTR Exons, 3’ UTR exons and Intergenic regions; number of ChIP peaks is displayed in bold, and percentage in italic; **(B)** distribution of Zfp125 ChIP peaks in the promoter region from −5Kbp to +1bp; **(C)** Zfp125 consensus sequence: motif-1 **(D)** Zfp125 consensus sequence: motif-2; chip-seq was performed using 3 different IP samples vs Input sample; **(E)** heat map analysis of Zfp125 motifs1+2 coordinates (x-axis) vs. Zfp125 ChIP peak coordinates within promoters (y-axis); the % overlap of x-y coordinates is shown at the bottom right.

KRAB-ZFPs promote formation of heterochromatin by attracting co-repressors (Frietze et al., 2010; Seah et al., 2019; Sripathy et al., 2006). To test if this was the case with Zfp125, we used a published adult mouse liver dataset of the heterochromatin marker H3K9me3-ChIP-seq (Wang et al., 2019) and found a ~50% overlap with Zfp125 ChIP peaks (Fig. 4A). After gene annotation (GENECODE Genes; release M24), we found that ~51% of the named Zfp125 ChIP peak containing genes also contained H3K9me3 peaks (Fig. S5A; Table S6). This supports the association between Zfp125 and heterochromatin, likely explaining how Zfp125 represses gene transcription.

**Figure 4.**
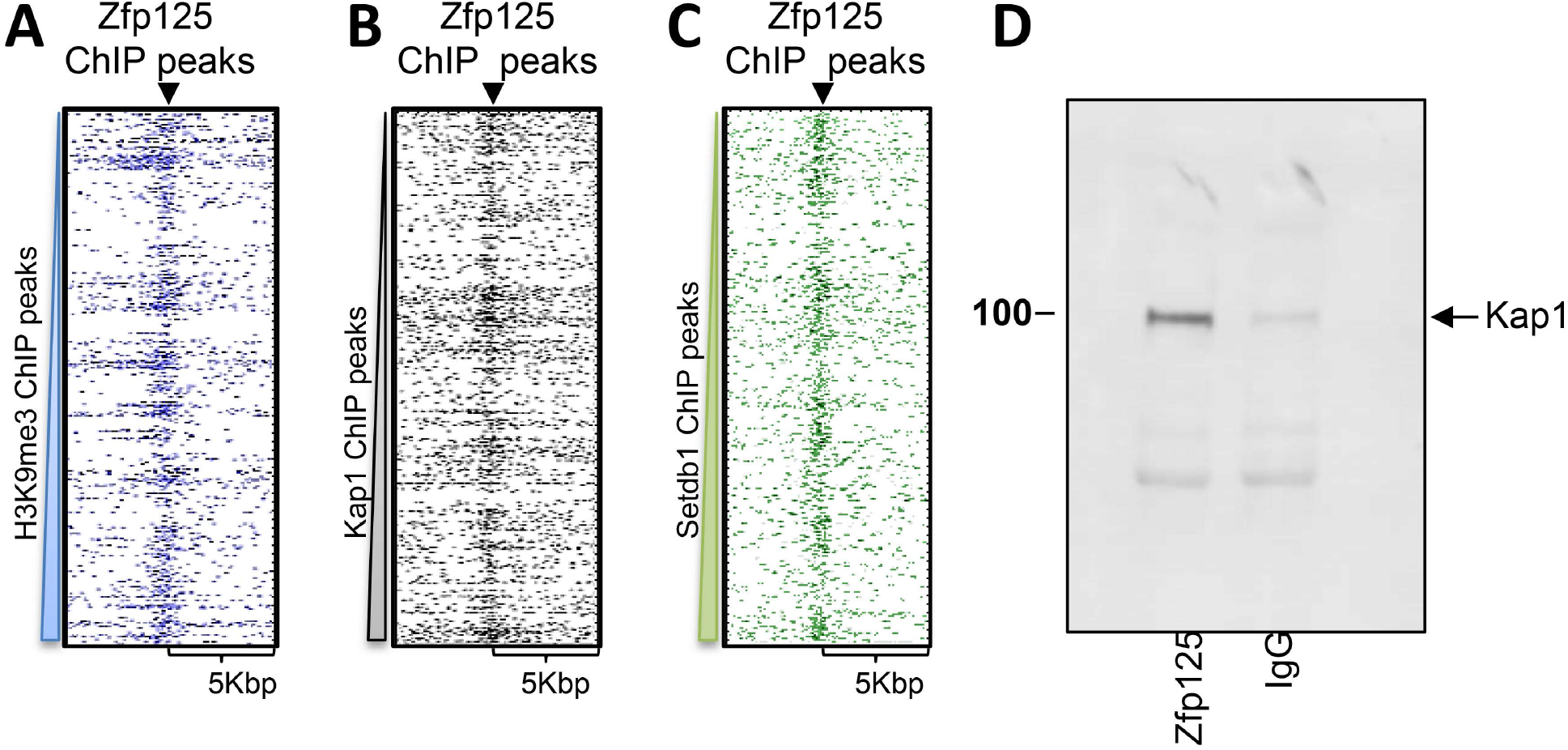
Mapping Zfp125 ChIP peaks. **(A)** heat map analysis of Zfp125 ChIP peak coordinates within promoters (x-axis) vs. H3K9me3 ChIP peak coordinates within promoters (y-axis) (Wang et al., 2019); **(B)** same as in **(A)** except that Kap1 ChIP peak coordinates within the promoters are on the y-axis (Kauzlaric et al., 2017); (**C**) same as in **(A)** except that Setdb1 ChIP peak coordinates within the promoters are on the y-axis (Kauzlaric et al., 2017); **(D)** cell lysates of AML12 cells stably expressing Zfp125 were immunoprecipitated with αZfp125 and pellets processed for western blot with αKap1.

Kap1 is one of the co-repressors that use the KRAB domain in ZFPs as a binding platform (Friedman et al., 1996; Kauzlaric et al., 2017). Thus, we used a published Hepa 1.6 liver cells dataset of Kap1-ChIP-seq (Kauzlaric et al., 2017) and build a promoter heat map of Zfp125 *vs* Kap1 ChIP peaks: ~10% of the Zfp125 ChIP peaks overlapped with Kap1 ChIP peaks (Fig. 4B). Using gene annotation, the overlap between Zfp125 ChIP peak-containing genes and Kap1 ChIP peak-containing genes was ~21% (Fig S6B; Table S5). Given this *in silico* evidence that Zfp125 can recruit Kap1, we next studied AML12 cells stably expressing Zfp125 and were able to pull down Kap1 using α-Zfp215 (Fig. 4D).

Setdb1 is a H3K9-specific histone-methyl-transferases (HMT) known to be recruited by Kap1 (O’Geen et al., 2007). To test if the Zfp125-Kap1 unit recruits Setdb1 to promote gene silencing, we used a published Hepa 1.6 liver cells dataset of Setdb1-ChIP-seq (Kauzlaric et al., 2017) and built a promoter heat map of Zfp125 *vs* Setdb1 ChIP peaks: ~20% of the Zfp125 ChIP peaks overlapped with Setdb1 ChIP peaks (Fig. 4C). The gene overlap analysis revealed that ~41% of the genes containing a Zfp125 ChIP peak also contained a Setdb1 ChIP peak (Fig S6C; Table S5). Altogether, these data indicate that the Zfp125-Kap1-Setdb1 repression unit functions to create heterochromatin and repress gene expression.

We also looked at the Zfp125 ChIP peaks located in intergenic regions, and asked whether they were near long-distance enhancer regions. A heatmap of intergenic Zfp125 coordinates *vs* the active enhancer marker H3K27ac or primed enhancer marker H3K4me1 ChIP peak coordinates revealed that the overlap was minimal (Praestholm et al., 2020)(Fig. S6D-G).

### The hepatic gene expression response to fasting is tailored by Zfp125

To define the mechanisms underlying the effects of Zfp125 on ketogenesis and gluconeogenesis, we performed microarray studies in liver of fed and fasted mice and compared the results with the Zfp125 ChIP-seq data.

In the fed state, liver-specific Zfp125 knockdown increased expression of 363 genes (>1.2-fold; p<0.05) (Table S6). An analysis of these genes revealed that 40 were metabolism-related (Table S7), including 22 genes involved in lipid metabolism, 11 in biological oxidation process, 12 in amino acid metabolism and transamination, and 3 involved in ketogenesis, i.e. Bdh1 and Bdh2 (that convert acetoacetate to BHB) and Hmgcs2 (catalyzes the first reaction of the ketogenesis pathway) (Fig. 5A-C).

**Figure 5.**
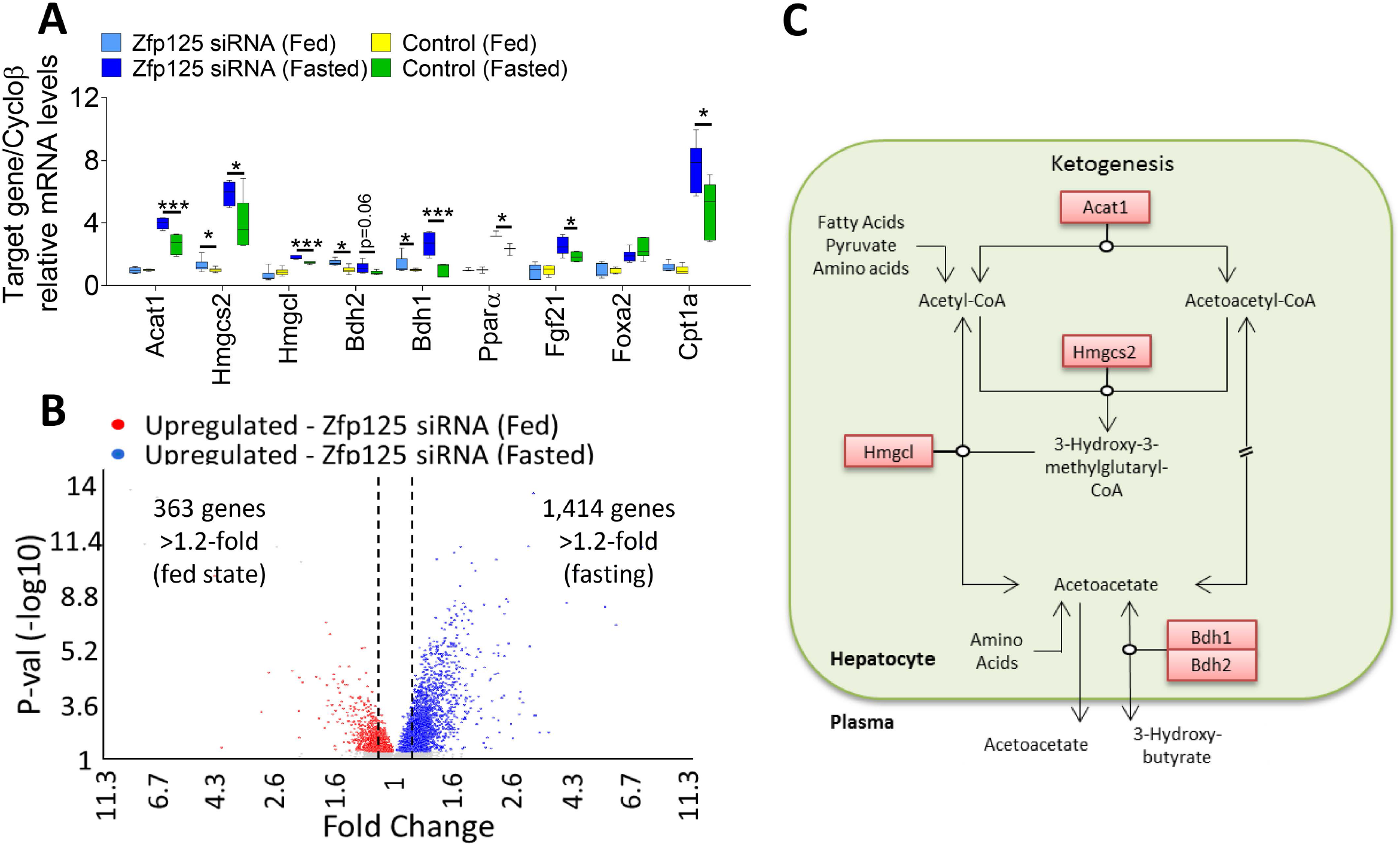
Effects of Zfp125 knockdown on hepatic gene expression. Mice received i.p. injections containing liposomes loaded with Zfp125 siRNA or non-target siRNA as in Fig. 1; some mice were fasted 36h immediately before being killed; livers were obtained and processed for RNA isolation; shown in **(A)** are the hepatic mRNA levels of selected as assessed by RT-qPCR; shown in **(B)** are the microarray results: a volcano plot of differentially expressed genes (>1.2-fold as indicated by doted lines; p<0.05) identified by Transcriptome Analysis Console (TAC); genes upregulated by Zfp125 siRNA in fed mice are shown in red and after 36h-fasting are shown in blue; **(C)** cartoon of hepatic ketogenesis pathway, highlighting genes (red box) upregulated by Zfp125 siRNA; values are the mean ± SEM of 6-12 independent samples. *P < 0.05, **P<0.01, ***P<0.001 as indicated; differences calculated by one-way ANOVA followed by Student-Newman-Keul’s test in A.

In the fasting state, there was a much more pronounced effect of liver Zfp125 knockdown on gene expression, enhancing the mRNA levels of 1,414 genes (>1.2-fold; p<0.05; Fig. 5A-B; Table S8). Among these, 129 genes were related to lipid metabolism, 31 were related to biological oxidation process, 70 involved with amino acids metabolism and transamination, 23 related with carbohydrate metabolism and 5 related to ketogenesis (Table S9 and Fig. 6). Some of the key genes involved included Bdh1, Bdh2, Hmgcs2 as well as Ppara and its downstream target Fgf21 that accelerates fatty acid catabolism, *Acat1 (*directs Acetyl CoA into the ketogenic pathway), Hmgcl (converts HMG-CoA to acetoacetate within the mitochondria) and Cpt1a (transports acylcarnitines into the mitochondrial matrix) (p<0.05; Fig. 5A-B and 6, Table S8 and S9).

**Figure 6.**
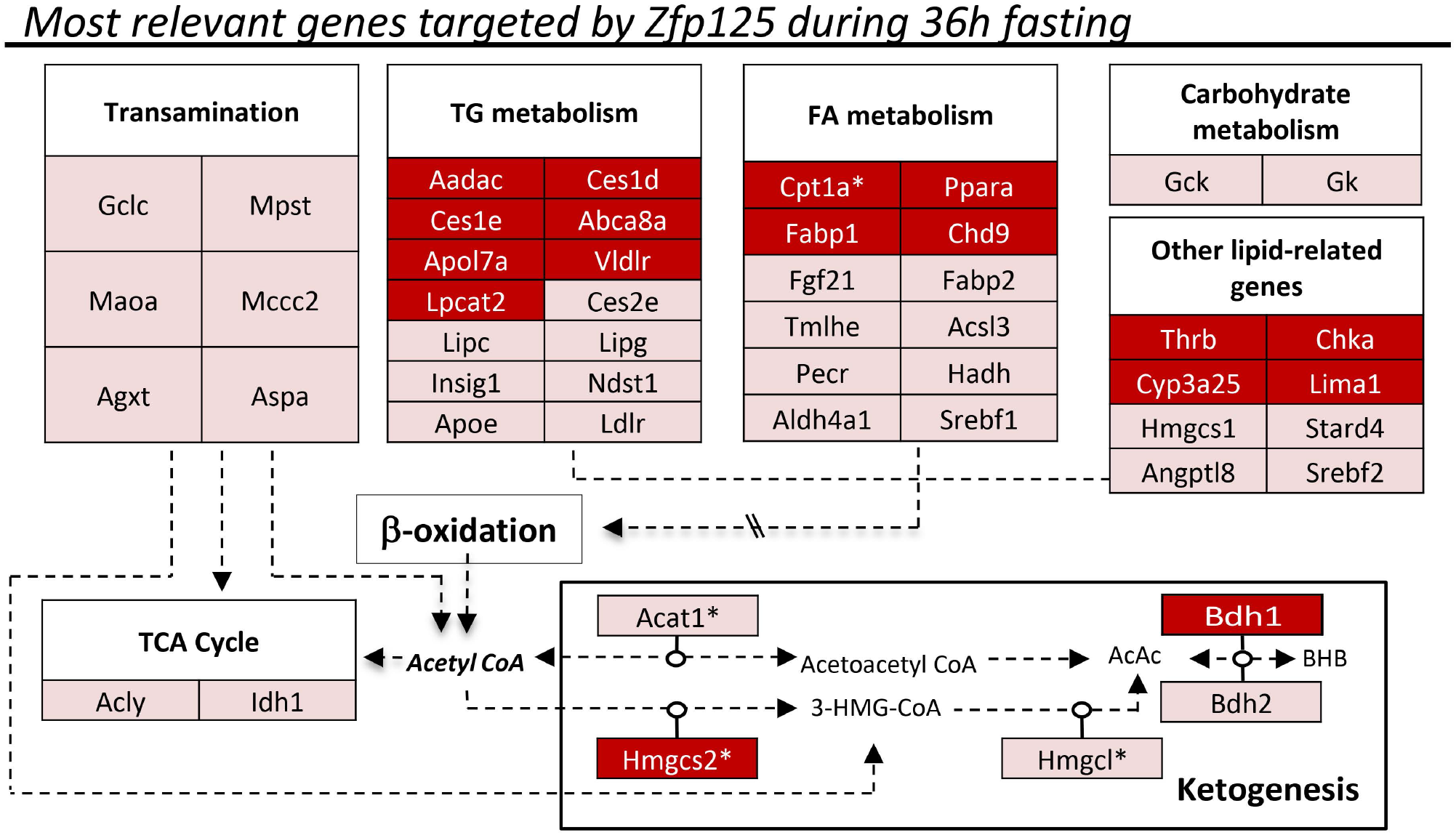
Zfp125 targets the promoter of key metabolic genes. **(A)** diagram (Partek-Flow platform) of the promoter regions of genes that (i) exhibit a Zfp125 ChIP peak (AML12 cells) and (ii) were up-regulated >1.2-fold in the liver of mice with liver-specific Zfp125 knockdown and fasted for 36h; in the diagrams, the blue areas indicate Zfp125 ChIP peaks and the red areas are peaks observed in the input samples; statistical difference between blue and red areas is indicated by the yellow bar; gene IDs are shown in the lower right corner of each box and TSS indicated by arrow; bp upstream of the TSS are indicated; **(B)** ChIP of AML12 cells stably expressing Zfp125 or empty vector with α-Zfp125 antibody followed by RT-qPCR targeted to the promoter of Hmgcs2; results are fold-enrichment vs. IgG antibody; **(C)** cartoon of mouse Bdh1 promoter, indicating two sites upstream of the TSS containing Zfp125 motif-2; **(D)** AML12 cells were transiently transfected with a luciferase reporter vector driven by the mouse Bdh1 promoter as shown in **(C)**; luciferase activity was normalized for β-galactosidase activity; values are the mean ± SEM of 3-12 independent samples;*P < 0.05, **P<0.01 as indicated as assessed through Student’s t-test in B and D.

### Mechanisms underlying transcriptional repression by Zfp125

An analysis of the ChIP-seq and microarray data revealed a number of genes that were potentially targeted by Zfp125, i.e. contained one or more Zfp125 ChIP peaks within their promoter regions and exhibited increased expression during Zfp125 knockdown (Fig. 6 and 7A; Table S10). For example, the presence of Zfp125 ChIP peak in the Hmgcs2 promoter was confirmed in AML12-Zfp125 cells using α-Zfp125 targeted to a region ~1600 bp upstream of the TSS of the Hmgcs2 promoter (Fig. 7B; Table S11). In the case of Bdh1, we also used a gene reporter system in which ~0.7kb of the mouse Bdh1 promoter drives a luciferase reporter (pGl3.Bdh1; Fig. 7C), having found a ~70% inhibition in AML12-Zfp125 (Fig. 7D).

**Figure 7.**
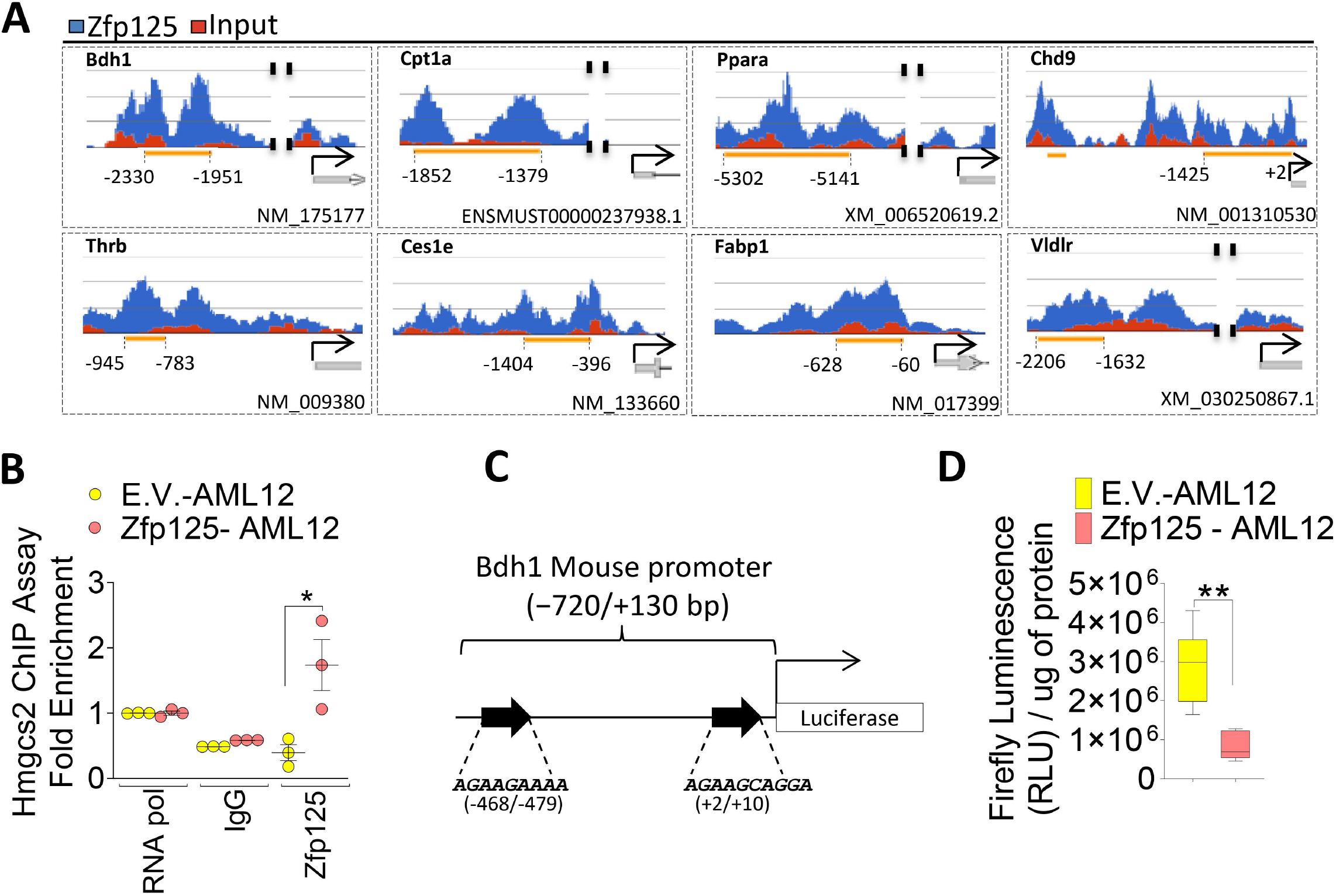
Relevant genes upregulated by Zfp125 siRNA in the mouse liver. Genes are grouped by metabolic pathways, as indicated. Genes identified by the microarray analysis and ChIP-seq analysis are shown in the red boxes. Genes containing asterisk were identified by PCR analysis. All the other genes (pink boxes) were obtained in the microarray analysis using Transcriptome Analysis Console (TAC; Table S9).

To gain further insight into the mechanisms involved in Zfp125-mediated gene repression we performed a series of ChIP assays using mouse liver chromatin and antibodies against the proteins that potentially interact with Zfp125 (Fig. 6, 7A). The setting was the promoters of 2 key genes targeted by Zfp125, i.e. Cpt1a and Bdh1 (Fig. 7A, 8B, G). While a weak ChIP Zfp125 signal was detected in both promoters, knocking down Zfp125 mRNA levels (45-50%) with liver-specific siRNA, reduced slightly further the ChIP Zfp125 signal (Fig. 8B,G). Kap1 ChIP signal was also low in both promoters, and it was further reduced in the Bdh1 promoter after Zfp125 knockdown (Fig. 8C,H). These changes after Zfp125 knockdown were associated with robust reduction in the ChIP signal with αSetdb1 (Fig. 8D,I), likely explaining the drop in ChIP signal with αH3K9me3 in both promoters (Fig. 8E,J).

**Figure 8.**
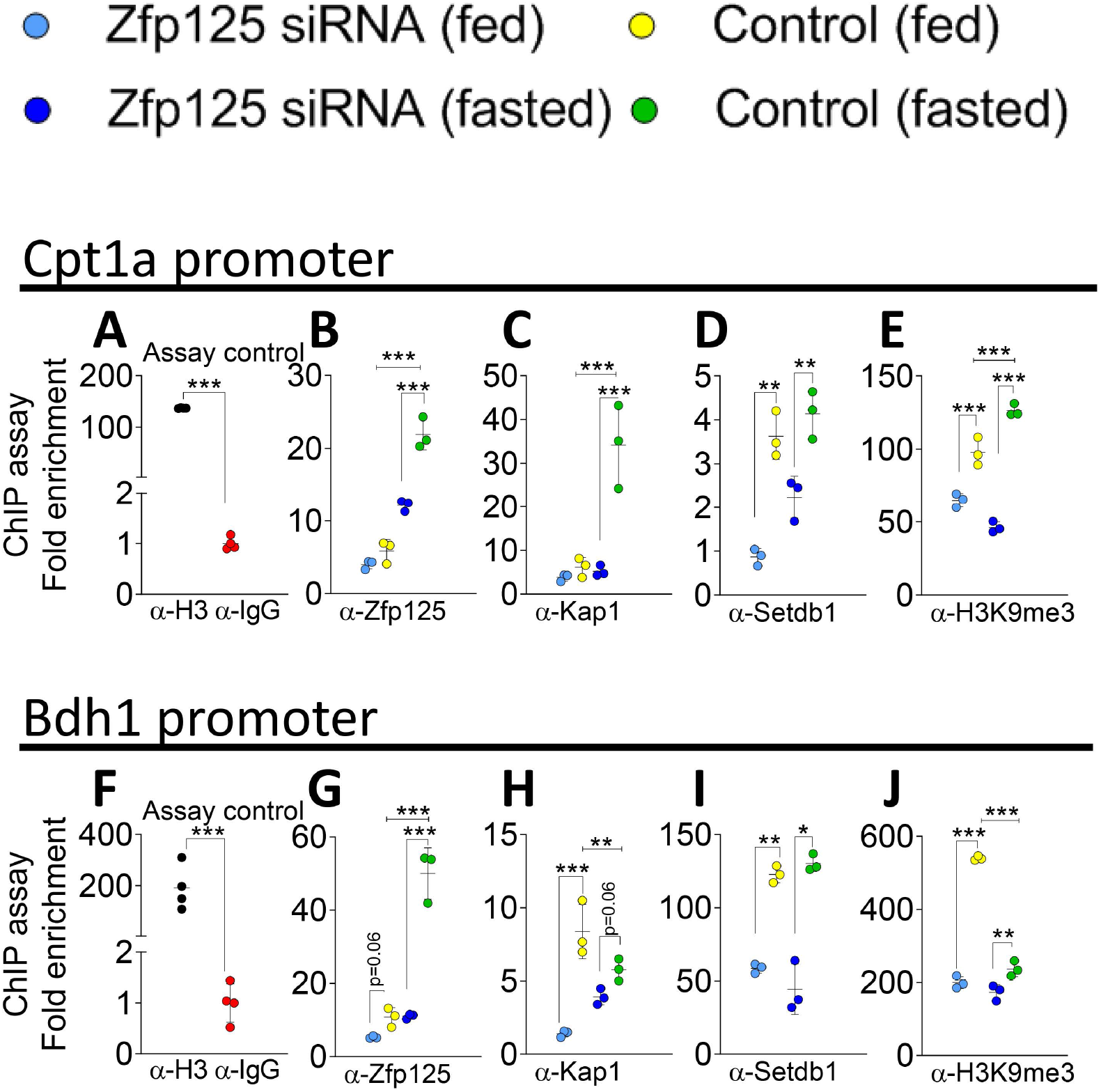
Transcriptional mechanisms initiated by Zfp125. **(A)** Positive and negative controls of liver ChIP assay respectively using α-Histone H3 or α-IgG followed by RT-qPCR targeted to the Cpt1a promoter. **(B)** Liver ChIP assay of fed or 36h-fasted mice using α-Zfp125, followed by RT-qPCR targeted to the promoter of Cpt1a; some mice had liver-specific Zfp125 siRNA or received scrambled RNA. **(C)** same as in **(B)**, except α-Kap1 was used. **(D)** same as in **(B)**, except that α-Setdb1 was used. **(E)** same as in **(B)**, except that α-H3K9me3 was used. **(F)** same as in **(A),** except that RT-qPCR was targeted to the Bdh1 promoter. **(G)** same as in **(B)** except that α-Zfp125 was used and RT-qPCR was targeted to the Bdh1 promoter. **(H)** same as in **(G)**, except that α-Kap1 was used. **(I)** same as in **(G)**, except that α-Setdb1 was used. **(J)** same as in **(G)**, except that α-H3K9me3 was used; values are the mean ± SD of 3-4 independent samples. *P < 0.05, **P<0.01, ***P<0.001 as indicated; differences calculated by one-way ANOVA followed by Student-Newman-Keul’s test.

There is activation of both Cpt1a and Bdh1 promoters during fasting, which also induces the expression of Zfp125 (Geisler et al., 2016; Kersten et al., 1999; Leone et al., 1999; Schlaepfer and Joshi, 2020). Thus, it makes sense that we saw a marked increase in the ChIP Zfp125 signal in both promoters when we used liver chromatin of mice that had been fasting for 36h (Fig. 8B,G). It is expected that these promoters are targeted by other transcription factors as well, hence the changes in ChIP signal of the corepressors during fasting were less consistent between both genes. For example, the increase in ChIP Kap1 signal was restricted to the Cpt1a promoter (Fig. 8C,H) and the ChIP Setdb1 signal was not at all affected by fasting in both promoters (Fig. 8D,I). In contrast, fasting increased the ChIP H3K9me3 signal in Cpt1a promoter (Fig. 8E) while reducing it in the Bdh1 promoter (Fig. 8J). Nonetheless, the effects of Zfp125 knock down were consistent across both promoters, i.e. a marked reduction in ChIP Zfp125 and Kap1 signals (Fig. 8B-C,G-H) as well as a drop in ChIP Setdb1 and H3K9me3 signals in the Cpt1a and Bdh1 promoters (Fig. 8D-E, I-J).

### Inhibition of ketogenesis by Zfp125 involves multiple steps

Hmgcs2 is the key rate limiting step in the ketogenesis pathway, converting acetyl-CoA into HMG-CoA (Shukla et al., 2017). The ~50% increase of Hmgcs2 mRNA levels caused by Zfp125 siRNA suggests direct inhibition of this step by Zfp125. This was tested by overexpression of HMGCS2 in AML12 cells stably expressing Zfp125. Indeed, HMGCS2 expression rescued BHB production, bringing the levels of BHB in the media to the levels observed in control cells (Fig. 9A). We next used a similar strategy and transiently expressed BDH1 in AML12-Zfp125 cells. BDH1 expression increased several-fold its mRNA (Fig. 9B) and accelerated accumulation of BHB in the medium by ~1.5-fold (Fig. 9B), completely rescuing the phenotype caused by Zfp125 expression.

**Figure 9.**
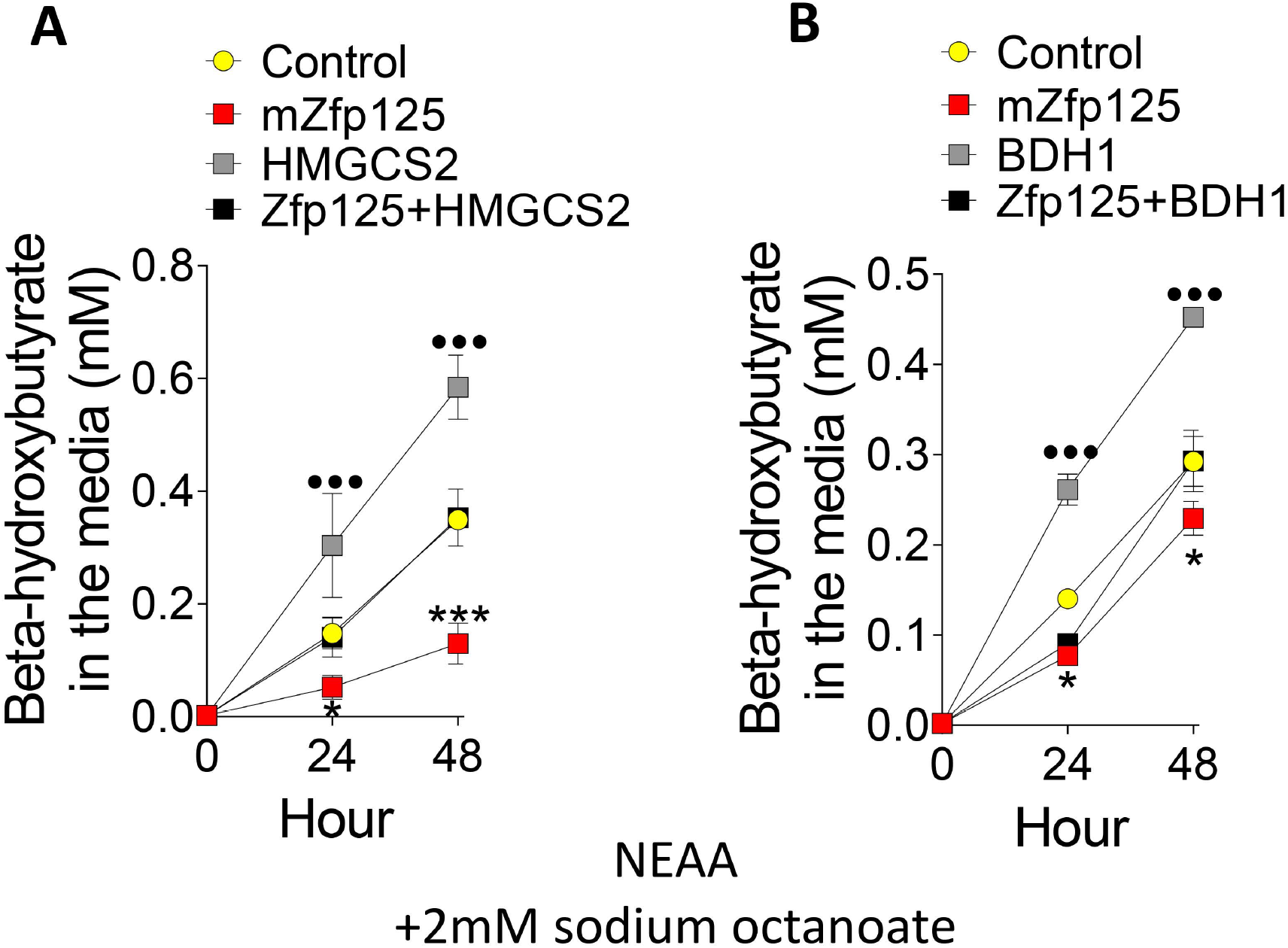
HMGCS2 and BDH1 are “at the center” of Zfp125 phenotype. **(A)** BHB levels in the medium of AML12 cells stably expressing Zfp125 or E.V. transiently transfected with human HMGCS2. **(B).** same as **(A**) except that cells were transiently transfected with BDH1 plasmid. All cells were incubated with medium containing nonessential amino acids (NEAA) and 2mM sodium octanoate. Values are the mean ± SD of 3-6 independent samples. *P < 0.05, ***P<0.001 vs. Control; ••• P<0.05 vs control. Differences calculated by one-way ANOVA followed by Student-Newman-Keul’s test in A-B.

### Zfp125 accelerates gluconeogenesis through allosteric stimulation of pyruvate carboxylase (PC)

The liver microarray and Zfp125 ChIP-seq studies did not identify genes or gene sets related to gluconeogenesis that were enriched or impoverished by Zfp125 knockdown (Table S6-9). Through RT-qPCR we also measured the four unique gluconeogenic hepatic enzymes, and identified three genes that were ~35-50% downregulated by Zfp125 knockdown, including G6pc, the enzyme that converts glucose 6-phosphate to glucose, and Pck1, the cytosolic rate-limiting enzyme that converts oxaloacetate to phosphoenolpyruvate (PEP) (Fig. 10A). We also saw that the cytosolic enzymes that produce oxaloacetate from malate/aspartate were inversely affected by Zfp125 knockdown, with Got1 mRNA levels being increased by ~60% and Me1 mRNA levels reduced by ~50% (borderline significant; Fig. 10A); no difference was observed in the mRNA levels of Fbp1, a gluconeogenic enzyme that catalyzes the hydrolysis of fructose 1,6-bisphosphate to fructose 6-phosphate or Pcx, the enzyme that catalyzes the carboxylation of pyruvate to form oxaloacetate (Fig. 10A).

**Figure 10.**
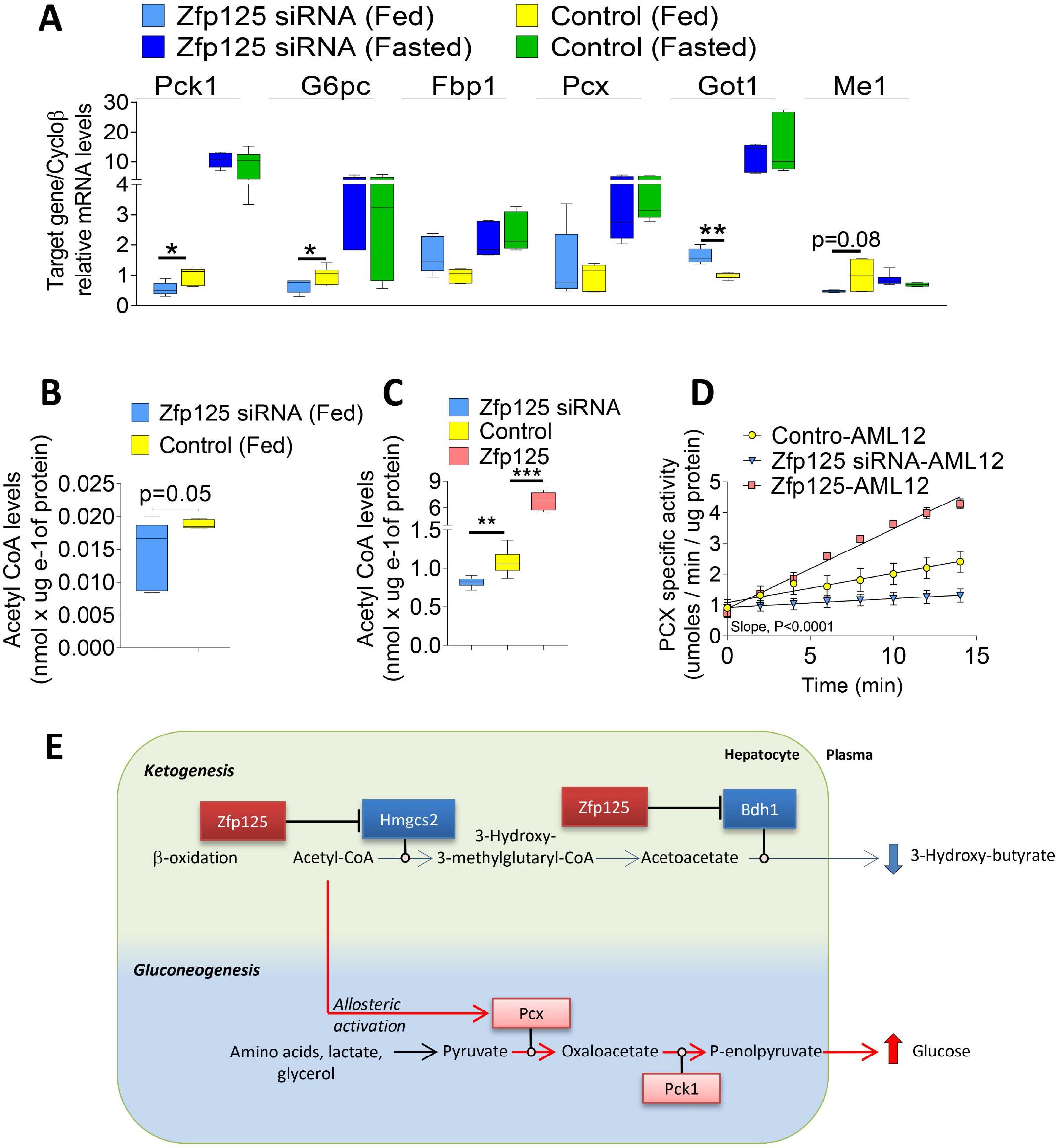
Zfp125 accelerates gluconeogenesis in liver and cells via Pyruvate Carboxylase activation. **(A)** mRNA levels of genes related to gluconeogenesis in the liver of fed/fasted mice transfected with Zfp125 siRNA or non-target siRNA, assessed by RT-qPCR. **(B)** Hepatic Acetyl CoA levels of mice fed *ad libitum.* **(C)** Acetyl CoA levels in AML12 cells stably expressing Zfp125, transiently transfected with Zfp125 siRNA or scrambled RNA. **(D)** Pyruvate carboxylase activity of cells in **(C)**; activity was measure over 15 min period. The difference between the slopes are P<0.0001. **(E)** Representative scheme of how Zfp125 sets the balance between hepatic ketogenesis and gluconeogenesis. Zfp125 represses key ketogenesis-related genes (as shown in blue boxes), reducing the output BHB. This leads to intracellular accumulation of acetyl-CoA, and activation of PC, resulting in increased gluconeogenesis. Values are the mean ± SEM of 5-18 independent samples.*P < 0.05, **P<0.01, ***P<0.001 as indicated; differences calculated by one-way ANOVA followed by Student-Newman-Keul’s test in A-C, and simple linear regression in D.

These findings made us look for alternate mechanisms to explain the Zfp125-induced changes in gluconeogenesis (Fig. 10B-E), including allosteric stimulation of PC by acetyl-CoA (Chourpiliadis and Mohiuddin, 2020; Jitrapakdee et al., 2008). PC converts pyruvate to oxaloacetate in the mitochondria, which is subsequently shuttled to the cytosol via the malate/aspartate mechanism. Thus, we tested whether inhibition of ketogenesis by Zfp125 could increase acetyl-CoA levels and accelerate PC activity. In mice, liver-specific Zfp125 knockdown reduced hepatic acetyl-CoA levels by ~50% (Fig. 10B), a change also seen in AML12 cells treated with Zfp125 siRNA (Fig. 10C). At the same time, acetyl-CoA levels were approximately 6-fold higher in AML12 cells stably expressing Zfp125 (Fig. 10C). This dramatic fluctuation in acetyl-CoA levels affected PC activity. In AML12 cells, Zfp125 siRNA slowed PC activity by ~50% while stable expression of Zfp125 accelerated PC activity by ~75% (Fig. 10D-E).

### Zfp125 minimizes induction of ketogenesis caused by glucagon or insulin resistance (IR), but gluconeogenesis is accelerated

We next studied the effects of Zfp125 on glucagon- and IR-induced ketogenesis and gluconeogenesis in 36h-primary cultures of mouse hepatocytes, knowing that glucagon increases Zfp125 mRNA by ~3.5-fold (Fig. 11A) and Zfp125 mRNA levels are ~1.6-fold higher in the liver of LIRKO mice, that have IR (Fernandes et al., 2018a). In this setting, Zfp125 expression was knocked down with siRNA by >75% or enhanced several-fold by transient expression of a Zfp125-expressing plasmid (Fig. 11B).

**Figure 11.**
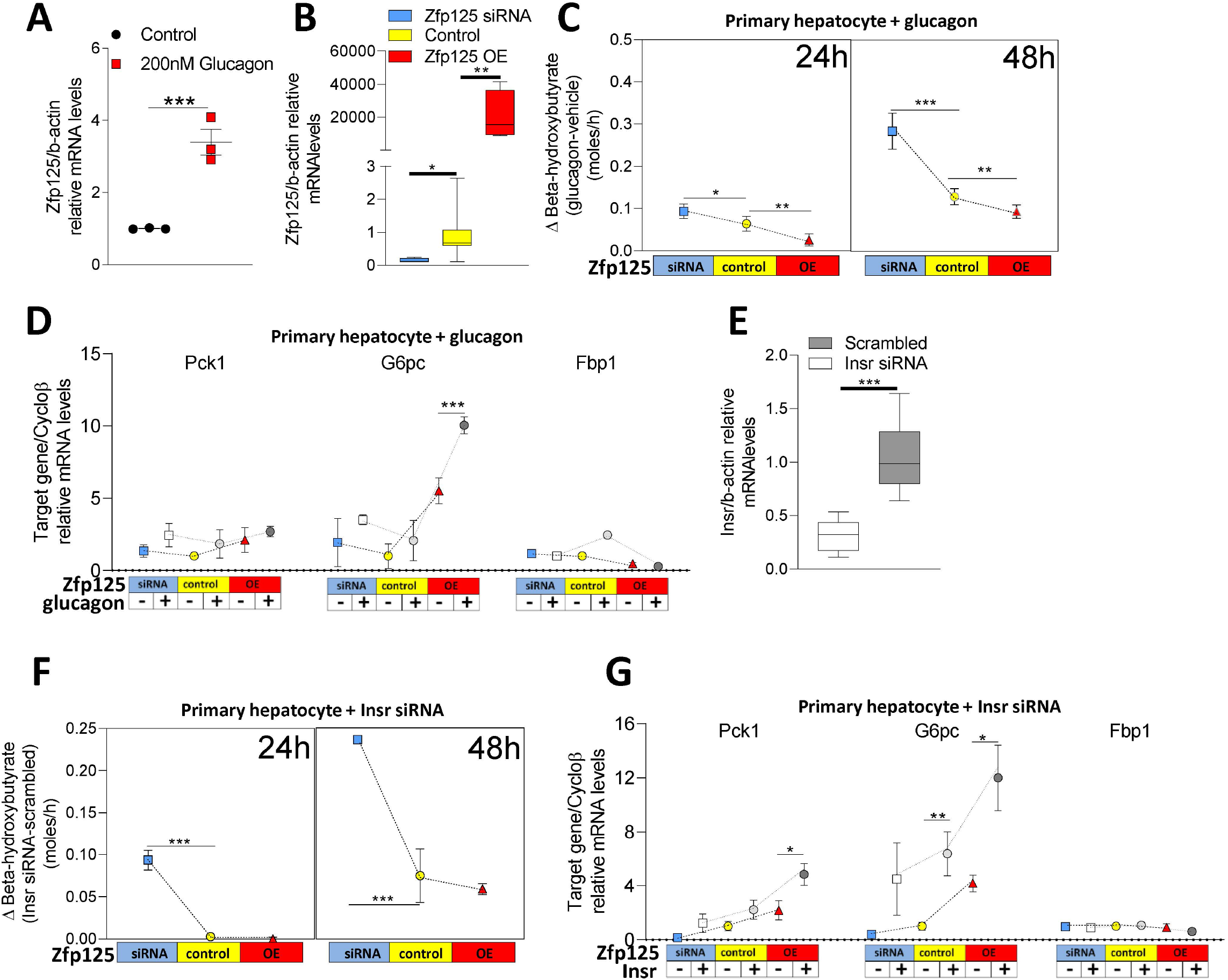
of Zfp125 on glucagon- and IR-induced ketogenesis and gluconeogenesis. **(A)** Zfp125 mRNA levels in primary hepatocyte cells treated with 200mM glucagon. **(B)** Zfp125 mRNA levels in primary hepatocyte cells overexpressing Zfp125, transiently transfected with Zfp125 siRNA or scrambled RNA;. **(C)** Delta BHB levels (gluca-gon-vehicle) in the medium of cells in **(B)** treated with or without 200nM glucagon; in all conditions medium were supplemented with non-essential amino acids and 2mM sodium octanoate for 48h; left box shows the levels after 24h and right after 48h. **(D)** Relative mRNA levels of gluconeogenic genes in cells showed in **(B)** and **(C)**. **E.** Insulin receptor (Insr) mRNA levels in primary hepatocyte cells transiently transfected with Insr siRNA or scrambled RNA. **F.** Delta BHB levels (Insr siRNA – Scrambled) in the medium of cells in of cells in **(B)** transiently transfected with Insr siRNA or scrambled RNA; in all conditions medium were supplemented with 20nM insulin, non-essential amino acids and 2mM sodium octanoate for 48h; left box shows the levels after 24h and right after 48h. **(G).** Relative mRNA levels of gluconeogenic genes in cells showed in **(D)** and **(F)**. Values are the mean ± SEM of 3-12 independent samples. *P < 0.05, **P<0.01, ***P<0.001 as indicated; differences were calculated using Student’s t-test in A and E, and one-way ANOVA followed by Student-Newman-Keul’s test in B, C-D, F-G.

Incubation of 36h-primary hepatocytes in medium containing NEAA and 2mM sodium octanoate with 200nM glucagon increased BHB production by ~150 umoles/h (Fig. 11C; Fig. S7A), whereas in the cells treated with Zfp125 siRNA the induction reached values ~300 umoles/h higher (Fig. 11C; Fig. S7A). Zfp125 overexpression only minimally reduced BHB production in response to glucagon, down to ~100 umoles/h (Fig. 11C; Fig. S7A). To assess gluconeogenesis in this setting, we looked at the expression of three key genes, i.e. Pck1, G6pc and Fbp1, but found that glucagon failed to have marked effects (Fig. 11D). Nonetheless, Zfp125 overexpression doubled the glucagon-induced G6pc, whereas it did not affect Pck1 and Fbp1 (Fig. 11D).

We knocked down insulin receptor (Insr) with siRNA in 36h-primary hepatocytes to promote IR in these cells, which reduced Insr mRNA levels by ~60% (Fig. 11E). Cells were kept in medium containing NEAA and 2mM sodium octanoate and 20nM insulin. IR increased BHB production in control cells by ~0.10 mmoles/h (Fig.11F; Fig. S7B), but in cells with Zfp125 knockdown IR-induced BHB production increased by ~0.25 mmoles/h (Fig. 11F; Fig. S7B). Overexpressing Zfp125 had little effect on IR-induced BHB production (Fig. 11F; Fig. S78). We also looked at the gluconeogenic genes and found that IR increased the expression of G6pc, but not Pck1 or Fbp1 (Fig. 11G). Whereas Zfp125 knockdown had little effect on IR-induction of these genes, Zfp125 overexpression amplified the IR induction of Pck1 and G6pc (Fig. 11G).

## Discussion

The key finding that liver-specific Zfp125 knockdown caused a rapid drop in RQ (Fig. 1A) led to the discovery that Zfp125 inhibits BHB production (Fig. 1C), while accelerating hepatic gluconeogenesis and HGP, both in mice and in two cell models (Fig. 1D, E, 2D). Zfp125 was found in the nucleus of hepatocytes (Fig. S5) and its ChIP peaks were distributed throughout the genome of AML12 cells (Fig. 3). However, the fact that no substantial overlap was observed between Zfp125 ChIP peak coordinates and the ChIP peaks of two enhancer markers H3K27ac/H3K4me1 (Fig. S6D-G), suggests that Zfp125 does not act remotely via distant enhancers. In fact, thousands of Zfp125 ChIP peaks were found in gene promoters (Fig. 3A). The coordinate overlap between Zfp125 and Kap1 ChIP peaks, as well as the co-immunoprecipitation of Zfp125 and Kap1, indicate that Zfp125 represses gene transcription by recruiting Kap1 to specific to the promoter of these genes (Fig. 4B, D).

Kap1 acts as a scaffold for a silencing complex that may contain the histone methyltransferase Setdb11 (Schultz et al., 2002), the nucleosome remodeling and deacetylation (NuRD) complex (Schultz et al., 2001), heterochromatin protein 1 (Hp1) (Sripathy et al., 2006) and DNA methyltransferases (Quenneville et al., 2012). While there is universal consensus that KRAB-ZFP-KAP1 induces formation of H3K9me3, only in developing cells does ZFP-mediated recruitment of Kap1 lead to DNA methylation (Quenneville et al., 2012; Seah et al., 2019). In differentiated cells, ZFP-Kap1-induced H3K9me3 formation allows for dynamic regulation of gene expression without involving DNA methylation (Quenneville et al., 2012). Such differences suggest that the composition of molecular complex recruited by Kap1-ZFP varies according to the developmental setting of the cell. Indeed, in the present studies we saw a marked overlap between Zfp125 and H3K9me3 ChIP peaks (Fig. 4A), indicating the involvement of a histone methyl transferase, possibly Setdb1, given the overlap between Zfp125 and Setdb1 ChIP peaks (Fig. 4C). This was confirmed with a series of ChIP assays targeted to the Cpt1a and Bdh1 promoters (Fig. 8). Indeed, binding of Kap1 to both of these promoters is dependent upon the presence of Zfp125 (Fig. 8B-C, G-H). In addition, knockdown of Zfp125 also disrupted recruitment of Setdb1 and diminished the amount of H3K9me3 in both promoters (Fig. 8D-E, I-J).

Repression of gene expression is at the center of the metabolic roles played by Zfp125. By directly repressing key genes involved in TG metabolism, fatty acid oxidation and ketogenesis (Fig. 6, Table S10), Zfp125 slows down production of BHB (Fig. 1C, 2B-C) and accelerates HGP (Fig. 1E, 2D). Zfp125 mRNA levels are increased in response to fasting via Foxo1 (Fernandes et al., 2018b) and glucagon (Fig. 11A), tailoring the hepatic transcriptional response to food deprivation (Fig. 5–7). While inhibition of ketogenesis seems to be purely transcriptional, the effect on gluconeogenesis involve transcriptional (i.e. Pck1 and G6pc; Fig. 10A, 11D, E) and non-transcriptional mechanisms via acetyl CoA-mediated allosteric activation of PC (Fig. 10B-C). These data are reminiscent of metabolic roles played by other KRAB-ZFPs (Ecco et al., 2017). For example, ZNF224 was shown to interact with KAP1 (Medugno et al., 2005) and repress the expression of human mitochondrial citrate carrier (Iacobazzi et al., 2009) and aldolase-A genes (Lupo et al., 2011). At the same time, overexpression of Zfp69 led to increased liver fat content and plasma TG levels in mice kept on HFD, along with downregulation of genes involved in glucose and lipid metabolism (Chung et al., 2015).

The present studies also pointed towards Zfp125 having a broader inhibitory role on hepatic gene expression. In the fed state, liver-specific Zfp125 knockdown increased the expression of hundreds of hepatic genes (Fig. 5B, Table S6) whereas during fasting that number increased to well over a thousand genes (Fig. 5B, Table S8). Based on the ChIP-seq data, only a fraction of these genes is likely to be directly inhibited by Zfp125, i.e. 43 genes in the fed state and 135 genes during fasting (Fig. 6, Table S10). Nonetheless, in each of the lipid-related areas, TG and fatty acid metabolism, β-oxidation and ketogenesis, there were key genes targeted by Zfp125 (Fig. 6, Table S10). It is striking that both Ppara and its co-activator Chd9, and its downstream target Fgf21, were up-regulated after Zfp125 knockdown (Fig. 5A, Table S8), suggesting strong inhibition of the Ppara pathway by Zfp125. This is particularly evident in β-oxidation and ketogenesis, where key genes such as Cpt1a, Hmgcs2 and Bdh1 were inhibited by Zfp125 (Fig. 5A, 6).

Hmgcs2 and Bdh1 play complementary roles in ketogenesis (Laffel, 1999) and were found at the center of the Zfp125-mediated mechanisms in the liver (Fig. 6, 7, 9A-B). Zfp125 ChIP peaks were present in the promoter of both genes (Fig. 7A-B), and both are upregulated by Zfp125 knockdown (Fig. 5A). Furthermore, the “Zfp125 phenotype” was rescued by overexpression of HMGCS2 or BDH1, bringing the BHB levels similar to those observed in control cells (Fig. 9A-B). This is reminiscent of the observation that mice treated with Hmgcs2 antisense oligonucleotide (Cotter et al., 2014; Geisler et al., 2019) as well as the Bdh1-deficient mice (Bdh1 KO) (Otsuka et al., 2020), also exhibit impaired synthesis of BHB. When fasted, these animals exhibited increased hepatic gluconeogenesis, mild hyperglycemia, and hepatic steatosis (Otsuka et al., 2020); all these features are caused by liver-specific Zfp125 expression (Fig. 1) (Fernandes et al., 2018b).

The fact that fasting-, glucagon- and IR-induced BHB production were all limited by Zfp125 indicates that Zfp125 fine tunes the rate at which gluconeogenesis and ketogenesis are accelerated during fasting, favoring the former over the latter (Fig. 1C-D; 2B-D; 11C, F). This bias towards gluconeogenesis suggests that Zfp125 could play a role in the excessive HGP observed in Type 2 Diabetes Mellitus (T2DM). In T2DM, increased rates of HGP result mainly from (i) excessive availability of gluconeogenic substrates; (ii) hepatic IR; and (iii) elevated glucagon levels that mimic hyperactivation of HGP seen during fasting (Sharabi et al., 2015). Hence, Zfp125 seems to be well positioned to influence HGP in T2DM: Zfp125 expression is induced by glucagon (Fig. 11A) and by Foxo1, is inhibited by insulin, and is elevated in the liver of the LIRKO mouse that has IR (Fernandes et al., 2018b).

The main Zfp125-dependent drive to accelerate gluconeogenesis was not transcriptional (Fig. 10) but rather the increase in acetyl-CoA levels and allosteric activation of PC activity (Fig. 10D). The increase in acetyl-CoA levels is likely to result from direct inhibition of BHB production, with a net positive balance of acetyl-CoA (Fig. 10B-E). The reduction in hepatic expression of two key gluconeogenic enzymes, Pck1 and G6pc, by Zfp125 knockdown (Fig. 10A) is also likely to contribute, but it was limited, most likely because these genes were not directly regulated by Zfp125. Nonetheless, Zfp125 did amplify the induction of both genes by glucagon and by IR (Fig. 11D, G). Thus, Zfp125 actions triggered a concerted effort to accelerate PC activity and the expression of key gluconeogenic genes, accelerating gluconeogenesis and HGP.

The present studies show that Zfp125 represses gene expression by recruiting Kap1 and Setdb1, and accelerating formation of heterochromatin. By repressing key metabolic genes, Zfp125 plays a physiological role in glucose homeostasis, tailoring the hepatic response to fasting, limiting production of BHB while accelerating gluconeogenesis and HGP. The fact that Zfp125 mRNA is increased during IR, and that it limits BHB production while inducing gluconeogenic genes suggests that Zfp125 might play a role in the excessive HGP and in the limited production of ketonic bodies observed in patients with T2DM. Defects in the regulation of gluconeogenesis can cause hepatic IR in obese nondiabetic and T2DM humans, which is aggravated by persistently high levels of glucagon (Sharabi et al., 2015). In these situations, insulin loses its ability to suppress gluconeogenesis, further increasing glucose concentration in the circulation (Sharabi et al., 2015).

## Methods

All experiments were planned and approved by the Institutional Animal Care and Use Committee at University of Chicago and Rush University Medical Center as described previously (Fernandes et al., 2018a; Fonseca et al., 2015).

### DNA Constructs and AML12-cell culture

AML12 mouse hepatocytes (American Type Culture Collection) were cultured in DMEM/F-12 medium (Gibco), and supplemented with 10% Fetal bovine serum (FBS), 5 ug/ml insulin, 5 ug/ml transferrin, 5 ng/ml selenium (ITS; Gibco) and 40 ng/ml dexamethasone (DEX) (Sigma). The full-length sequence of mouse Zfp125 (mZfp125; Fig. S3A and S4A) was initially amplified by PCR and syn-thetized by the GeneScript Company. The cloning was done using a pUC57 vector, with additional 5’ sequence (gctagc), additional 3’ sequence (gaattc), EcoRV blunt cloning site and a LAC promotor. In order to create clones that stably express Zfp125, Zfp125 product was subcloned into a pCI-Neo mammalian expressing vector (Promega), and construct was transfected into AML12 cells. Stable cells expressing Zfp125 were generated using G418 (400ug/mL) for 7 days. The functionality of this plasmid was confirmed by qRT-PCR. The mZfp125 is ~ 74% homologous to ZNF670, the corresponding gene in humans (hZfp125) (Fernandes et al., 2018a). For the experiments using hZfp125, *Zfp125* siRNA and controls (Empty vector and Non-target siRNA), cells were generated and cultured as described previously (Fernandes et al., 2018a). The mouse *Bdh1* promoter from −720 bp to +130 bp subcloned into the pGL3-basic luciferase vector (Promega) was kindly given by Dr. Hitoshi Shimano and Dr. Yoshimi Nakagawa from University of Tsukuba (Nakagawa et al., 2016). The cDNAs from the human *BDH1* (cloneID HsCD00356339) and the human *HMGCS2* (cloneID HsCD00641671) inserted in the pANT7_cGST plasmids were obtained from the Arizona State University Biodesing DNASU Plasmid Repository. Human BDH1 or HMGCS2 plasmids were transiently transfected in AML12 cells stably expressing Zfp125 or controls. After 24h of transfection complete medium was added and cells were prepared for assay. Insulin Receptor siRNA was obtained by GE Healthcare Dharmacon, Inc (ON-TARGETplus Mouse Insr siRNA; L-043748-00-0010). All transfections were done using Lipofectamine 2000.

### Perfusion and primary hepatocyte culture

Primary hepatocytes were obtained from C57B6 mice. Initially, Perfusion medium (GIBCO) was thawed 30 min before digestion. Mice were anesthetized with ketamine and xylazine, and needle was placed in portal vein. Perfusion medium was done at a rate of 4.5 ml/minutes during 5-6 minutes. Subsequently, liver digest medium (GIBCO) was used at the same rate during 6-8 minutes. Liver was excised and placed in a 100mm plate filled with cold wash medium (DMEM:F12 medium) and teared into pieces. Pieces were purified using a 100μm filter and centrifuged at 50xg for 3 minutes. Pellet of cells were re-suspended using 40% cold percoll medium (ThermoFisher) and centrifuged at 150xg for 7 minutes. Pellet of cells were 2 times washed with cold wash medium followed by centrifugation at 50xg for 3 minutes. Cells were re-suspended and cultured using collagen coated plates. Complete DMEM:F12 medium, supplemented with 10%FBS, 5 ug/ml insulin, 5 ug/ml transferrin, 5 ng/ml selenium (ITS; Gibco) and 40 ng/ml dexamethasone (DEX) (Sigma) were added 5h after isolation. In some experiments, 5h-cultured hepatocytes were transfected with Zfp125 plasmid, Zfp125 siRNA, Insr siRNA and controls (Empty vector and Non-target siRNA) using Lipofectamine 2000 in a 0.1% FBS medium. After 5h complete medium (DMEM:F12) was added, and after 19h cells were prepared for assay.

### Immunofluorescence in primary hepatocyte culture

Primary hepatocytes obtained from C57b6 mice were fixed with 4% paraformaldehyde, permeabilized with 0.1% Triton X-100 in PBS and incubated in normal goat serum (5 %) in PBS to block non-specific antibody binding sites. For immunofluorescence labeling, cells were incubated with a rabbit polyclonal antibody for Zfp125 (1:400; ThermoFisher, Table S13). The secondary antibody was an anti-rabbit DyLight 488 (1:200; Vector, Table S13). Cell nuclei were stained with 4’,6-diamidino-2-phenylindole (DAPI; 1:10,000; ThermoFisher, Table S13). Confocal images were acquired using an inverted laser scanning confocal microscope (Nikon Eclipse C1). Catalog number for all antibodies is shown in Table S13.

### Immunoprecipitation (IP) studies in AML12

AML12 stably expressing hZfp125 were used for IP studies using the Immunopreciptation Kit from Abcam. Cells pellet were resuspended in Lysis Buffer (non-denaturing) with a Protease Inhibitor Cocktail (Abcam), sonicated and incubated at 4°C for 30 minutes. Cell homogenates were centrifuged for 10 minutes at 10000 *g at* 4°C and transfer to fresh tubes. The samples were quantified by Bradford, and 250-400 μg of total protein were used per IP. In some samples, pellets were incubated with mouse α-Flag (20 μg; Sigma-Aldrich, Table S13) or mouse α-IgG (1 μg; Millipore, Table S13) overnight at 4°C. For the Co-IP experiment, pellets were incubated with rabbit α-Zfp125 (2 μg) or rabbit α-IgG (2 μg; Cell Signaling, Table S13) overnight at 4°C. The following day 40 μl of protein A/G Sepharose were added in each sample and incubated at 4°C for 1 h. After washing with 1 ml of wash buffer three times, the beads were eluted with 40 μl of SDS-sample buffer and boiled during 5 minutes for protein denaturation and Western blot analysis. Catalog number for all antibodies is shown in Table S13.

### Western blot studies

Western blot analyses utilized 30-60 μg of total protein or immunoprecipitate pellets. Proteins were resolved on a 4%–12% SDS-PAGE gel and transferred to a PVDF membrane (Immobilon-FL, Millipore), incubated with indicated antibodies overnight at 4°C and subsequently quantitated with the LiCOR Odyssey instrument with Odyssey Image Studio software using 2 different infrared channels. The following antibodies were used at indicated dilutions: 1:5000 for α-Zfp125 (Table S13), as previously described (Fernandes et al., 2018a); 1:500 α-Kap1 (Abcam; Table S13); and 1:1000, Lamin A/C (Cell Signaling, Table S13). Catalog number for all antibodies is shown in Table S13.

### Chromatin immunoprecipitation (ChIP) assay in AML12

In AML12 cells stably expressing hZfp125, ChIP assays were performed using an EZ-Chip Kit (Millipore). Chromatin lysates were prepared, pre-cleared with Protein-A/G agarose beads, and immune precipitated with antibodies against the *Zfp125* (Fernandes et al., 2018a), RNA pol II (Millipore; Table S13), or normal mouse IgG (Millipore, Table S13) as previously described (Fernandes et al., 2018a). Beads were extensively washed before reverse crosslinking. DNA was purified and subsequently analyzed by RT-qPCR using primers flanking the proximal binding sites located in the mouse *Hmgcs2* promoter.

### ChIP-sequencing (seq) in AML12

ChIP was initially performed in AML12 cells stably expressing hZfp125, using the SimpleChIP^®^ Plus Enzymatic Chromatin IP Kit (Magnetic Beads; Cell Signaling). Chromatin lysates were prepared, pre-cleared with Protein G magnetic beads, and immunoprecipitated with antibodies against the Zfp125 (in house; Table S13) as previously described (Fernandes et al., 2018a). Beads were extensively washed before reverse crosslinking. Chip-enriched DNA was purified and 10ng of fragmented input DNA was used for the generation of the library using the TruSeq Chip Sample Prep Kit (Illumina). Briefly, Chip-DNA was quantified using a High sensitivity DNA kit Qubit ds DNA HS Assay Kit. The ends of the DNA fragments were repaired using the AMPure XP Beads. A single ‘A’ nucleotide was added to the 3’ ends of the blunt fragments to prevent them from ligating to one another during the adapter ligation reaction. Multiple indexing adapters were used to ligate the ends of the DNA fragments, preparing the samples for hybridizations onto a flow cell. Ligation products were them purified and amplified, thought PCR reaction. Finally, library was validated and sequenced using an Illumina High Throughput Sequencing PE100. Fastq files obtained from the sequencing were aligned to the mouse assembly *Mus musculus* – mm10 using the Partek-flow platform. Peak calling was done with MACS2 analysis with criteria of p<0.05, and a MFold rang of 3-30. Peaks were annotated with the mouse genome GENECODE GENE (release M24), and genes associated with the annotated peaks were filtered based on regions next to the Transcriptional Start Site. Finally, gene set enrichment, pathway enrichment, and de novo motif analysis were performed in the list of these genes. Heat maps analysis were generated using Easeq software. The following published ChIP-seq datasets were used: GSM2339529 (SETDB1 ChIP-Seq in Hepa 1.6 cells; Mus musculus; ChIP-Seq); GSM2339528 (KAP1 ChIP-Seq inHepa 1.6 cells; Mus musculus; ChIP-Seq); GSM2339536 (Total input Hepa 1.6 cells; Mus musculus; ChIP-Seq)(Kauzlaric et al., 2017); GSM3753284 (baseline WT H3K9me3);

GSM3753283 (input WT H3K9me3) (Wang et al., 2019); GSM3679089 (WT H3K27Ac) and GSM3679121 (WT H3K4me1) (Praestholm et al., 2020).After re-analyzing all ChIP-seqs using the same parameters as Zfp125 ChIP-seq, BED files were generated and used for heat map studies.

### Luciferase gene reporter assay in AML12

AML12 stably expressing Zfp125 cells or EV were transfected with pGL3Basic mouse Bdh1 promoter as previously described (Fernandes et al., 2018a). 24h later cells were processed for luciferase activity using Dual-Luciferase Reporter (DLRTM) Assay System (Promega).

### Ketone body output assay

To measure ketone bodies output in cells, primary hepatocytes and AML12 cells were pre-washed with DBPS and incubated with a 1:1 mixture of DMEM/F12 no phenol red (Gibco) with SILAC Advanced DMEM/F-12 Flex Media containing Non-essential amino acids. Unless indicated otherwise, all cells were treated with 2mM sodium octanoate. Medium samples were collected at 0h, 24h and 48h after the incubation. BHB levels in the medium were determined using the β-OH butyrate colorimetric assay kit (Abcam). In some sets of experiment, 20nM insulin or 200nM glucagon was supplemented in the 1:1 mix of medium every 24h. Ketone body levels were corrected by protein levels of cells.

### Glucose output assay

To measure glucose output in cells, cells were initially prewashed with DPBS and pre-incubated with SILAC Advanced DMEM/F-12 Flex Media, no glucose, no phenol red (Gibco) during 3h, for depletion of glycogen. After incubation, cells were washed with DPBS and treated with 2mM sodium pyruvate in SILAC Advanced DMEM/F-12 Flex Media, no glucose, no phenol red (Gibco) during 3h. Medium samples were collected at1h and 3h after the incubation. Glucose level in the medium was determined using the Glucose Colorimetric/Fluorometric assay kit (Abcam).

### Pyruvate Carboxylase assay in AML12

Pyruvate carboxylase activity was measured in AML12 cells transfected with Zfp125 siRNA, stably expressing mZfp125 or controls as previously described (Payne and Morris, 1969). Briefly, cells pellet were resuspended in 100 mM tris°HCl buffer (pH 8.0) and broken with a tissue tearor (Biospec Products,Inc). Cells were centrifuged to remove cell debris and kept on ice. Two cocktails (MilliQ H2O, tris HCl, NaHCO3, MgCl2, Acetyl CoA, DTNB, Citrate Synthase and ATP) with or without pyruvate were prepared and kept on ice. After 10 minutes to allow the temperature of the solutions to equilibrate, cell extracts were simultaneously added to the cocktail and the mix was counted during 15 min at a wavelength of 412nm. Calculation of activity was obtained by calculating the Total volume used × Dilution of the cell extract ×[Rate]experimental – [Rate]control×1000/ Volume of cell extract used× Molar extinction coefficient for reduced DTNB× Concentration Factor of cell extract.

### Animal Experiments

Unless indicated otherwise, all animals were 8-week old male C57BL/6J mice that were kept at room temperature (22°C), with a 12h dark/light cycle and were fed *ad libitum* with a standard chow diet and water. All animals were euthanized by asphyxiation in a CO_2_ chamber. Tissues and plasma were rapidly collected and frozen in liquid nitrogen and stored at −80 °C until analysis.

### Liposome-mediated liver-specific transfection

*In vivo* transfections were performed with a liver transfection kit (Altogen Biosystems), as previously described (Fernandes et al., 2018a). Zfp125 plasmid, Zfp125 siRNAs were transfected into male C57BL/6J mice according to the manufacturer’s protocol, via intraperitoneal injection. Briefly, 120 μg of DNA or siRNA diluted in 100μl of RNase-/DNase-free water was incubated with 50μl of Transfection reagent-1 (Altogen) for 15 min at room temperature. Transfection Enhancer Reagent-2 (Altogen) was added to the tubes containing DNA/siRNA+Transfection re-agent-1 and incubated for 5 min at room temperature. After incubation, sterile solution containing 5% glucose (w/v) was added to the mixture according to table in the manufacturer’s protocol. Transfectant+DNA/siRNA mixture were injected in mice at 0h, 24h, 48h, 72h, 96h, and killed at the 120h time point.

### Studies in the Comprehensive Lab Animal Monitoring System (CLAMS)

Animals were admitted to a comprehensive laboratory animal monitoring system (CLAMS; Columbus Instruments), kept at 22 °C, and studied as described previously (Fonseca et al., 2015). Animals were placed in the CLAMS with free access to food and water, allowing them to acclimatize in individual metabolic cages for 48h. Subsequently, metabolic profiles were generated in successive 26-min cycles during 120h. Respiratory quotient (RQ) and food consumption were monitored continuously.

### Fasting experiment

Age and sex matched mice were randomly divided into *ad libitum* chow fed or fasting groups, with only access to water. The fasting period was 36 h from 6:00am – 06:00pm. Mice were subsequently sacrificed by asphyxiation with CO2, and plasma was collected directly from the heart, and stored on ice with EDTA to prevent clotting. Blood was then centrifuged at 8,000 x g for 8 min to separate cells and plasma. Livers were frozen in liquid nitrogen and stored at −80°C for later analysis.

### Pyruvate tolerance test (PTT)

Mice transiently transfected with mZfp125, Zfp125 siRNA or scrambled RNA were prepared for PTT. Experiment were performed in mice fasted for 12 hours followed by intraperitoneal injection of pyruvate (2 g/kg body weight) dissolved in saline solution. Blood glucose was measured using a One-Touch Ultra glucometer (LifeScan) with a single drop of tail blood at T= 0, 15, 30, 60, 90, and 120 minutes after challenge.

### Chromatin immunoprecipitation (ChIP) assay in liver

Using liver of mice transiently transfected with Zfp125 siRNA or scrambled RNA, ChIP assays were performed using the SimpleChIP^®^ Plus Enzymatic Chromatin IP Kit (Magnetic Beads; Cell Signaling). Chromatin lysates were prepared, pre-cleared with Protein G magnetic beads, and immunoprecipitated with antibodies against Zfp125 (ThermoFisher; Table S13), Kap1 (Abcam; Table S13), Setdb1 (ThermoFisher; Table S13) and H3K9me3 (Abcam; Table S13). For control samples, liver was immunoprecipitated using α-Histone H3 (Cell Signaling; Table S13) or α-IgG (Cell Signaling; Table S13). DNA was purified and subsequently analyzed by RT-qPCR using primers flanking the proximal binding sites located in the Ppara, Cpt1a and Bdh1 promoters. Catalog number for all antibodies is shown in Table S13.

### Microarray Analysis

RNA obtained from liver of mice transfected with *Zfp125* siRNA or controls, fed or 36h-fasted, was extracted with the RNeasy Mini Kit (Qiagen) and processed for microarray at the core facility in the University of Chicago (Chicago, IL). Gene expression was evaluated using Mouse Gene 2.0 ST arrays (Affymetrix, Inc., Santa Clara, CA). Gene expression data were preprocessed using Affymetrix Expression Console. Differential expression analysis was performed in Affymetrix Transcriptome Analysis Console (TAC) to identify individual genes.

### Biochemical analyses

Blood was collected into EDTA tubes and plasma was separated by centrifugation. Glucose levels in the plasma or cell medium were measured using the Glucose Colorimetric/Fluorometric assay kit (Abcam). β-hydroxybutyrate levels in the plasma or medium were measured using the beta Hydroxybutyrate Colorimetric assay Kit. Acetoacetate levels in the plasma or medium were measured using Acetoacetate Colorimetric assay Kit. Plasma free fatty acid level was assessed by Colorimetric/Fluorometric assay kit (Abcam). Plasma insulin level was obtained using Insulin (Mouse) ELISA Kit (Abnova). Acetyl Co-A levels in the liver and cells were obtained using the Acetyl-CoA Fluorometric Assay Kit (Biovision).

### Liver Histology

After liver extraction and dissection, part of the liver tissue was immersed in buffered formalin (10%) and fixed for 24h. Paraffin-embedded tissues were sectioned and processed as described (Fernandes et al., 2014; Ueta et al., 2012) for staining with periodic acid-Schiff (PAS; to indicate glycogen content) (17,18). For immunohistochemistry analysis, liver samples were initially frozen in OCT and prepared by the Integrated Light Microscopy Core Facility from The University of Chicago. To detect Zfp125 expression, liver section was incubated with α-Zfp125 using a 1: 2500 dilution.

### Gene expression analysis

RNA was obtained as described previously (Fonseca et al., 2015). Specific mRNA levels were quantified by RT-qPCR (StepOnePlus real time PCR system, Applied Bioscience) using SYBR Green Supermix (Quanta Biosciences). Standard curves consisting of 4-5 points of serially diluted mixed experimental and control group cDNA were included and the coefficient of correlation was consistently >0.98, with an amplification efficiency of 80-110%. Table of primers is shown at Table S12.

### Statistics

All data points are mean ± SEM and prepared using PRISM (GraphPad Software, San Diego, CA). One-way ANOVA was used to compare more than two groups, followed by the Student-Newman-Keuls’ test to detect differences between groups. The Student’s t-test was used when only two groups were part of the experiment; p<0.05 was used to reject the null hypothesis.

## Acknowledgement

We thank Integrated Light Microscopy Core Facility from The University of Chicago, especially Shirley Bond and Terri Li, for their assistance with histology images. This work was supported by NIDDK (R01 DK77148 – ACB) and National Research, Development and Innovation Office (K-125247 – BG). The authors are also grateful to Drs. Hitoshi Shimano and Yoshimi Nakagawa from University of Tsukuba, Japan for providing the Bdh1 plasmid.

## Author Contributions

G.W.F. and B.M.L.C.B. performed experiments, analyzed data and prepared figures; T.L.F., assisted with the CLAMS analysis. F.S.L. performed immunofluorescence; O.N. and S.C.R. assisted in the data analysis of the microarray; G,B. prepared plasmids and discussed/revised the manuscript; I.C.K discussed/revised the manuscript; A.C.B. planned and directed all studies and prepared the manuscript.

## Legend to Supplemental Figures

**Figure S1. (A)** Experiment design performed in adult mice admitted to the CLAMS; after acclimatization, mice received i.p. injections of liposomes directed to the liver loaded with Zfp125 siRNA at 0h, 24h, 48h, 72h and 96h; control animals received liposomes loaded with non-target siRNA; some animals were fasted for 36h immediately before killing; all animals were killed at 120h after the first injection and processed for: **(B-E)** immunohistochemistry of liver sections; α-Zfp125 (1:2500) was used; magnification is indicated. **(F)** relative mRNA levels of Zfp125 in the soleus muscle, as assessed by RT q-PCR. **(G)** Relative mRNA levels of Zfp125 in eWAT of mice shown in **(A)**, as assessed by RT q-PCR. **(H).** Delta body weight of mice shown in **(A)**. **(I)** Liver weight of mice shown in **(A)**. **(J)** eWAT weight of mice shown in **(A)**. **(K)** Lean carcass (skeletal muscle and bones) weight of mice shown in **(A)**. **(L)** Food intake of mice shown in **(A)**. Values are the mean ± SEM of 6 independent samples; differences were calculated using Student’s t-test.

**Figure S2. (A)** Fluctuation of respiratory quotient (RQ) of mice shown in Fig. S1A; some animals were fasted for 36h immediately before killing (Fig.S2A); image shows the final 60h of the experiment. **(B-E)** Representative liver section of mice in **(A);** periodic acid-schiff (PAS) was used to detect hepatic glycogen, as shown by arrows; magnification is indicated **(F)** Delta body weight of mice shown in **(A)**. **(G)** Liver weight of mice in **(A)**. **(H)** eWAT weight mice shown in **(A)**. **(I)** Lean carcass (skeletal muscle and bones) weight of mice shown in **(A).** Values are the mean ± SEM of 6-12 independent samples. *P < 0.05 vs. respective control; differences were calculated using Student’s t-test.

**Figure S3. (A)** Mouse Zfp125 gene and its coding sequence. The gene (light blue) is 47138 nucleotides long and is located in the chromosome 12 (20874125 to 20921262), and includes 7 exons and 6 introns (1). The promoter region contains 3 GC box and 1 CAAT box next to the transcriptional start site (TSS; as indicated). A Foxo1 binding site is also indicated in purple, approximately 2,500 bp upstream of the TSS. Coding sequence (2; red) starts in the 3^rd^ exon (methionine is indicated with an arrow) and ends with a TAA stop codon at the 6^th^ exon (indicated by arrow). There is a KRAB-A domain box in the N-terminal region, located in the 4^th^ exon, and 10 C2H2 domains towards its C-terminal region, entirely located in the 6^th^ exon. Using a 95% query cover, the human gene with the highest degree of homology to Zfp125 is ZNF670, exhibiting a ~74% homology, as shown in **B**. The human gene (light blue) is 44175 nucleotides long and is located in the chromosome 1 (247034637 to 247078811), including 4 exons and 3 introns (3). Coding sequence (4; orange) starts in the 1st exon (methionine is indicated with an arrow) and ends with a TAA stop codon at the 4^th^ exon (indicated by arrow). There is a KRAB-A domain box in its N-terminal region, located in the 2nd exon, and 10 C2H2 domains towards the C-terminal region, entirely located in the 4^th^ exon.

**Figure S4. (A)** Mouse Zfp125 nucleotide sequence (coding region). **(B)** Human Zfp125 nucleotide sequence (coding region).

**Figure S5. (A)** Representative Immunohistochemistry images of liver sections of mice shown in Fig. S1A with α-Zfp125 (1:2500); magnification is indicated. Zfp125 expression is present in the cytoplasm and in the nucleus (arrows). **(B)** Immunofluorescence images of cultured primary hepatocytes with α-Zfp125 and DAPI; arrows show nuclear expression. **(C)** Western blot of cytoplasm (left) and nuclei (right) fractions of AML12 cells stably expressing Zfp125 (lane 1) or Empty Vector (lane 2); lane 3 shows negative control (HEK293 cells); **(D)** Western blot with α-Zfp125 of α-Flag immunoprecipitated total lysates of AML12 cells stably expressing Zfp125; lane 1 is AML12 cells stably expressing Zfp125; lane 2 is AML12 cells stably expressing empty vector.

**Figure S6. (A)** Overlap analysis of genes with a Zfp125 peak within the promoter regions vs H3K9me3 ChIP-seq (Wang et al., 2019) (Table S5). **(B)** Same as (A) except that overlap was Zfp125 ChIP-seq vs the Kap1 ChIP-seq (Kauzlaric et al., 2017) (Table S5). **(C)** Same as (A) except that overlap was Zfp125 ChIP-seq vs Setdb1 ChIP-seq (Kauzlaric et al., 2017) (Table S5); the percentage overlap is shown; coordinates were annotated with GENECODE GENE (release M24). **(D)** Heat map analysis of Zfp125 ChIP peak coordinates within intergenic regions (x-axis) vs. H3K27ac ChIP peak coordinates within intergenic regions (y-axis) (Praestholm et al., 2020). **(E)** same as in **(D)** except that H3K4me1 ChIP peak coordinates within intergenic regions were used (Praestholm et al., 2020). **(F)** Heat map analysis of Zfp125 ChIP peak coordinates within promoter regions (x-axis) vs. H3K25ac ChIP peak coordinates within promoter regions (y-axis) (Praestholm et al., 2020). **(G)** same as in **(F)** except that H3K4me1 ChIP peak coordinates within promoter regions were used (Praestholm et al., 2020).

**Figure S7. (A)** BHB levels in the medium of primary hepatocytes transiently expressing Zfp125 (OE), Zfp125 siRNA, or scrambled RNA; cells were treated with 200mM glucagon or vehicle; medium was supplemented with non-essential amino acids and 2mM sodium octanoate; **(B)** same as (A) except that cells were also transiently transfected with Insr siRNA or non-target siRNA and were not treated with glucagon; 20nM insulin was used as indicated; values are the mean ± SEM of 3-12 independent samples. *P < 0.05, **P<0.01, ***P<0.001 as indicated; differences were calculated using Student’s t-test.

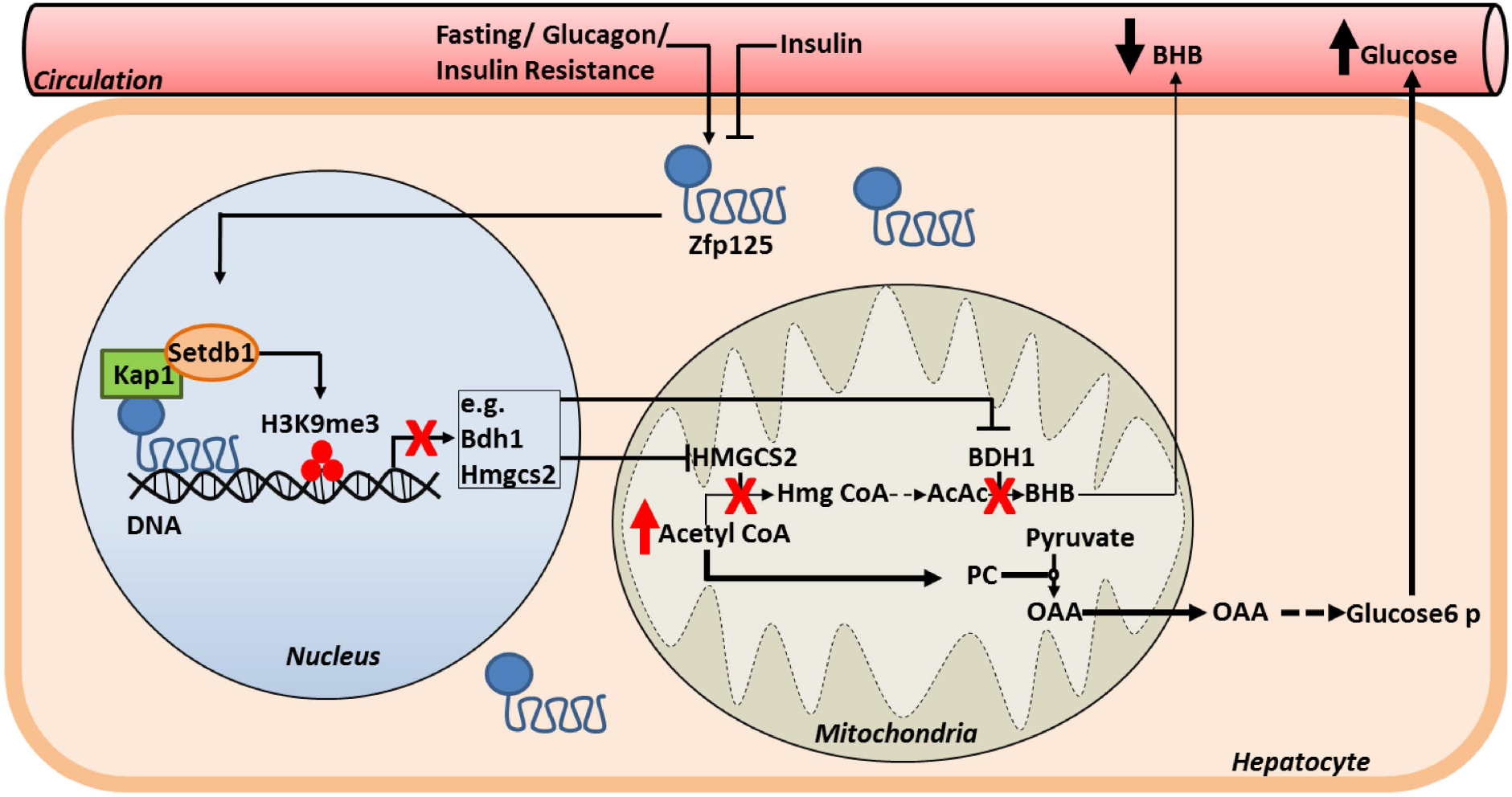
Graphical abstact

