## Supplemental Figures and Tables for "The transcriptional repressor Zfp125 modifies hepatic energy metabolism in response to fasting and insulin resistance": Fernandes et al - Supplemental Fig v4 - 2020.pdf

# A

### Design of the experiment

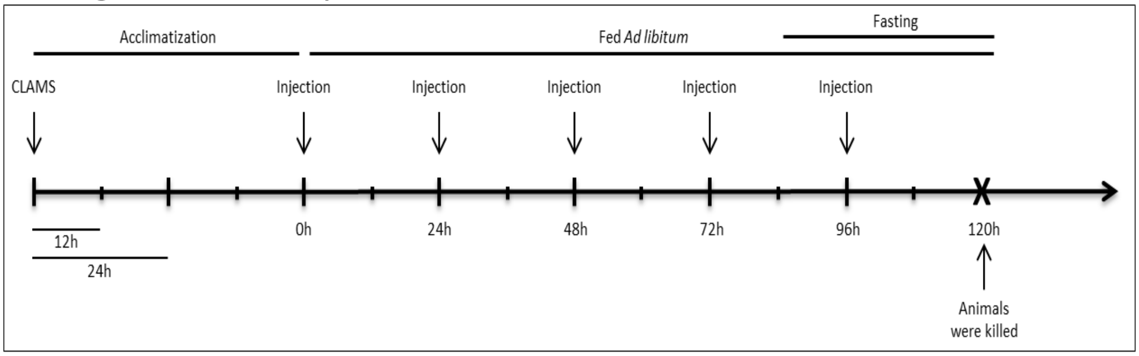


### Immunohistochemistry

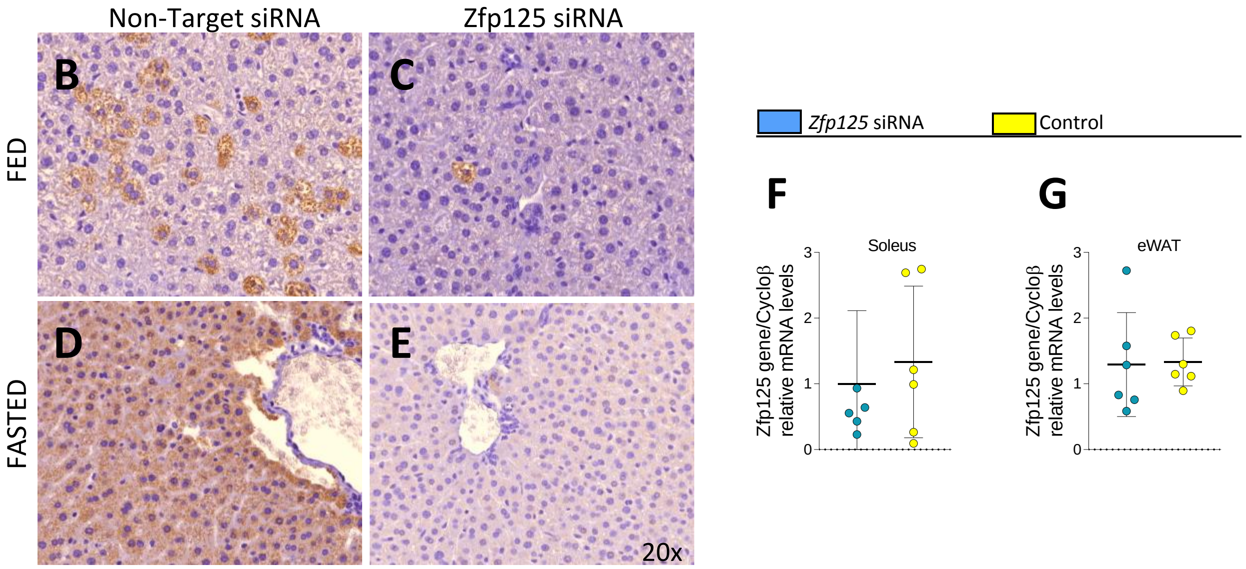


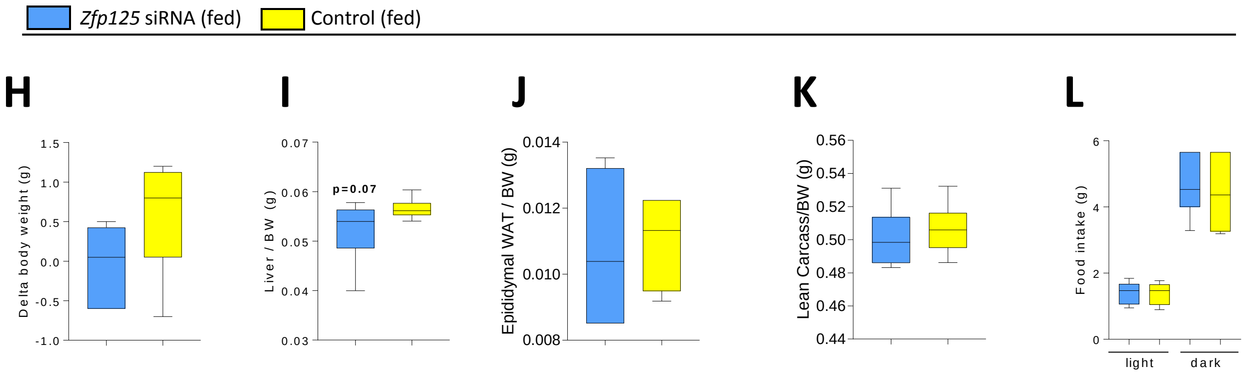


Figure S1

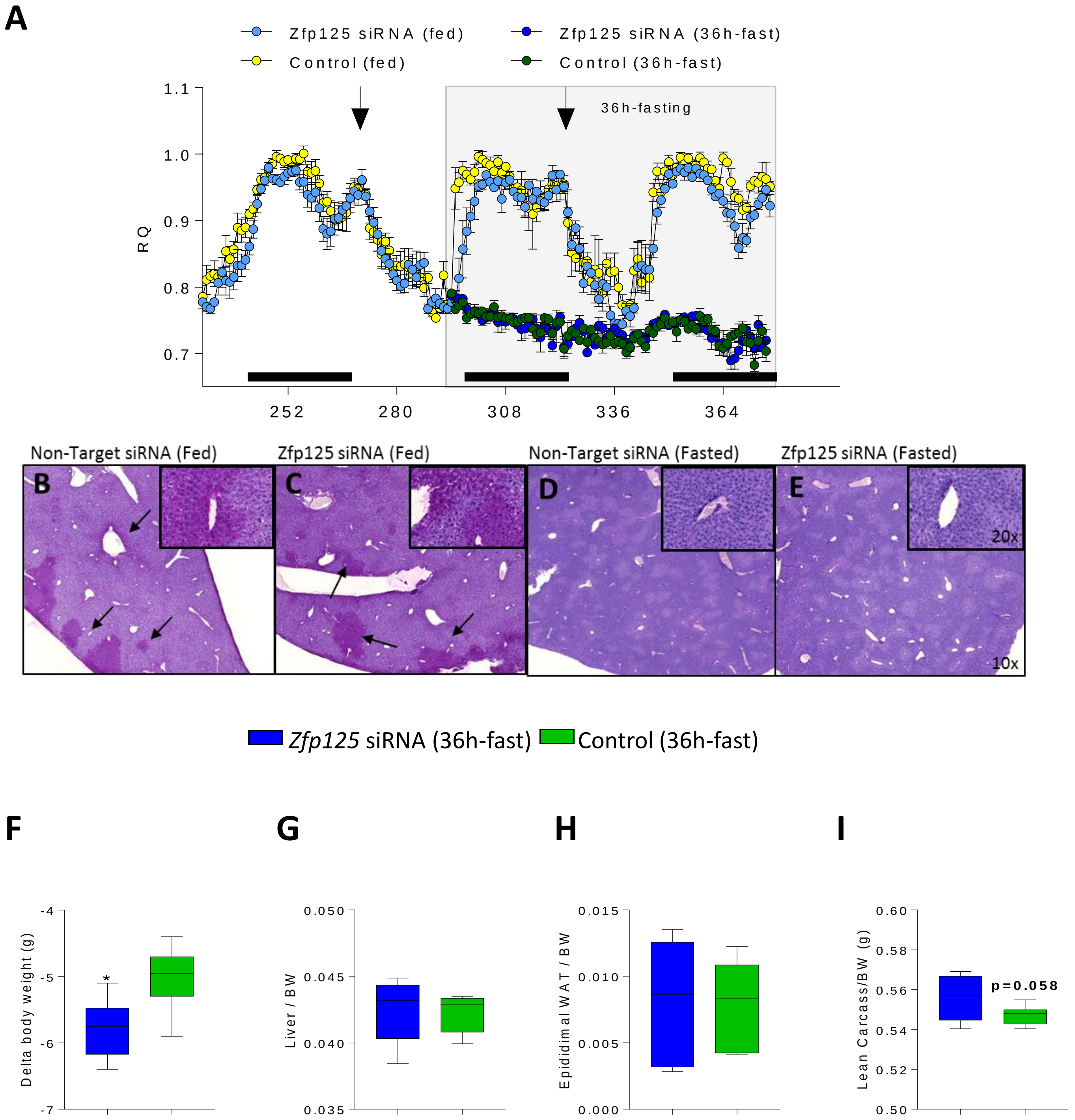
A
Zfp125 siRNA (fed)
Zfp125 siRNA (36h-fast)
Control (fed)
Control (36h-fast)
36h-fasting
RQ
1.1
1.0
0.9
0.8
0.7
252
280
308
336
364
Non-Target siRNA (Fed)
Zfp125 siRNA (Fed)
Non-Target siRNA (Fasted)
Zfp125 siRNA (Fasted)
B
C
D
E
20x
10x
Zfp125 siRNA (36h-fast)
Control (36h-fast)
F
G
H
I
Delta body weight (g)
Liver / BW
Epididimal WAT / BW
Lean Carcass/BW (g)
*
p=0.058

Figure S2

A.

Mouse Zfp125

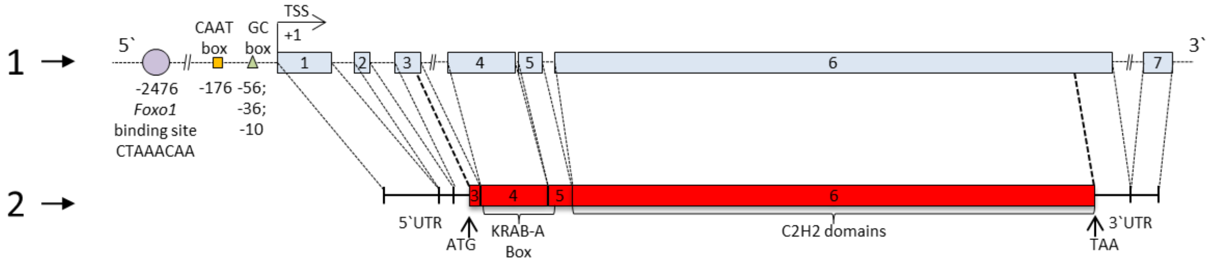


B.

Human Zfp125

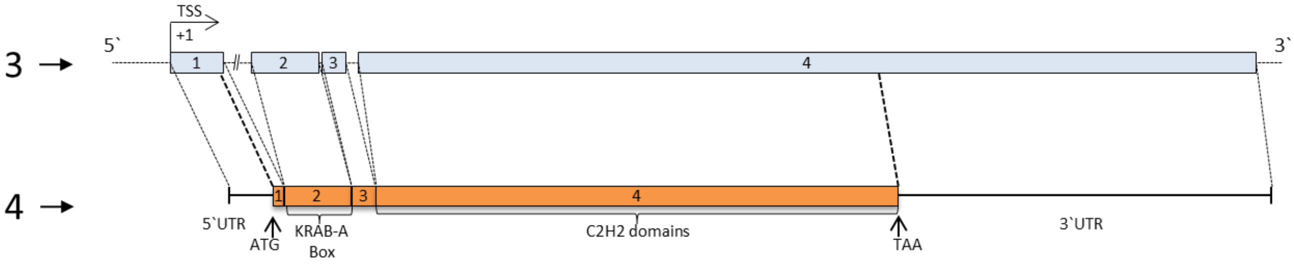


Figure S3

# A

**Zfp125 sequence**

ATGGATGCAGTGACTTATGATGATGTGCATGTGAACTTCACTGGGGAAGAGTGGAATTTGCTGGATCCTTCCCAGAAGAGTCT
CTACAAAGATGTGATGCTGGAGACCTACTGGAACCTCACTGTTATAGTTGAAGGACATTGTCAAAGTTCAAGAAGAAATGAAA
GGTATGTAAGTAATCATAGTGGAGAGAAACTCTATGAATATAATGAACGTTCTAAAGCCTTTTCATGTCCCAGTCATCTCCAAT
GTCATAAAAGAAGACAAATTGGAGAGAAAACACATGAACATAACCAATGTGGTAAAGCCTTTCCAACACCCAGTCATCTTCAA
TATCATAAAAGAACCCATACTGGAGAGAAACCTTATGAATGTCATCAATGTGGTCAAGCCTTTAAAAACTGCAGTCAACTCGA
ACATTATAAAAGAACACGTACTGGAGAGAAATCTTATGAATGTCGTCAATGTGGTAAAGCCTTTTCACAACACAGTCATCTCCA
ACGTCATAAAAGAATACATACTGGAGAGAAACCCTATGAGTGTAATCAATGTCGTAAAGCGTTTTCACAACACAGTCTTCTCCA
ACGTCATAAAAGAACACATACTGGAGAGAAACCCTATGAATGTAATCAATGTGGTAAAGTCTTTGCACATCTCAGTAGTTTAA
AATGTCATCACAGAACACATACTGGAGAGAAACCCTATGAATGTAATCAATGTGGTAAAGCCTTTGCAGGTCATAGTCATCTC
CAATGTCATAAAAGAACACATACTGGAGAGAAACCCTATGAATGTAATCAATGTGGTAAAGCCTTTGCACTACTCAGTAGTTTC
CAATGTCATAAAAGAACACATACTGGAGAGAAACCCTATGAATGTCATCAATGTGGTAAAGCCTTTGCACAACTCAGTAGTCT
CCACTGTCATAAAAGAACACATACTGGAGAGAAACCCTATAAATGTCATCAATGTGGTAAAGCCTTTTCACAACACAGTCATCT
TCAATATCATAAAAGAACACATACTGGAAAGAAACCCTAA

# B

**ZNF670 sequence**

ATGGATTCAGTGTCATTTGAAGATGTGGCTGTGGCCTTTACTCAGGAGGAGTGGGCTTTGCTGGATCCTTCTCAAAAGAAT
CTCTACAGAGATGTGATGCAAGAAATCTTCAGGAACCTGGCTTCTGTAGGAAACAAATCAGAAGACCAGAATATCCAAGA
TGACTTCAAAAATCCTGGGAGAAATCTAAGTCATGTGGTAGAGAGACTGTTTGAAATTAAAGAAGGCAGTCAATATGGAG
AAACCTTCAGCCAGGATTCAAATTTGAATCTGAATAAGAAAGTTTCTACTGGAGTAAAGCCATGTGAATGCAGTGTGTGTG
GAAAAGTCTTCATATGTCATTCAGCCCTTCATAGGCACATCCTGTCTCACATTGGAAACAAACTATTTGAGTGTGAGGAATG
TCCAGAGAAGTTATATCATTGCAAACAATGTGGGAAAGCCTTTATCTCTCTCACCAGTGTTGACAGACACATGGTAACACA
CACTAGTAATGGGCCTTATAAGGGTCCAGTGTATGAGAAGCCTTTTGATTTTCCTAGTGTATTTCAAATGCCTCAGAGCACT
TACACTGGAGAGAAAACATATAAATGTAAACATTGTGATAAAGCCTTCAATTATTCAAGTTATCTTCGTGAACATGAAAGA
ACTCATACTGGAGAGAAACCCTATGCATGTAAGAAATGTGGTAAATCATTCACTTTTTCCAGTTCTCTTCGCCAACATGAAA
GATCTCATACTGGAGAGAAACCCTATGAATGTAAGGAATGTGGCAAAGCCTTCAGTCGTTCCACTTACTTGGGAATACATG
AAAGAACGCATACTGGAGAAAAACCCTATGAATGTATAAAATGTGGCAAAGCCTTTAGATGTTCCAGAGTCCTCAGAGTCC
ATGAAAGGACTCACAGTGGAGAAAAGCCCTATGAATGTAAACAATGTGGTAAAGCCTTCAAATATTCTAGTAACCTATGTG
AGCATGAAAGAACTCACACTGGAGTGAAACCTTATGGATGTAAGGAATGTGGTAAGTCGTTTACTTCTTCCAGTGCCCTTC
GAAGCCATGAAAGGACTCATACTGGAGAAAAACCCTATGAATGTAAGAAATGTGGTAAAGCCTTCAGTTGTTCCAGTTCCC
TTCGAAAGCATGAAAGAGCTTATATGTGGTAA

Figure S4

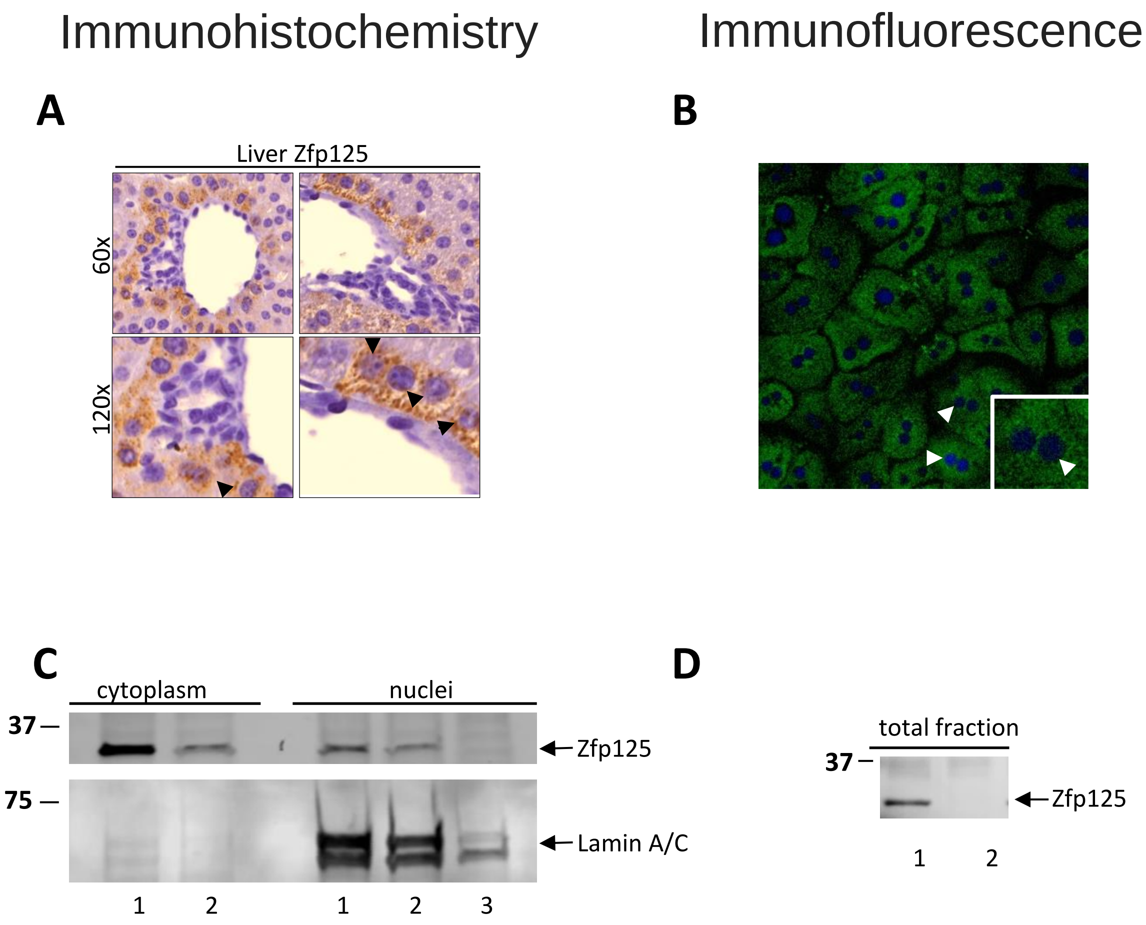
Immunohistochemistry
Immunofluorescence
A
Liver Zfp125
60x
120x
B
C
cytoplasm
nuclei
37
75
Zfp125
Lamin A/C
1
2
1
2
3
D
total fraction
37
Zfp125
1
2

Figure S5

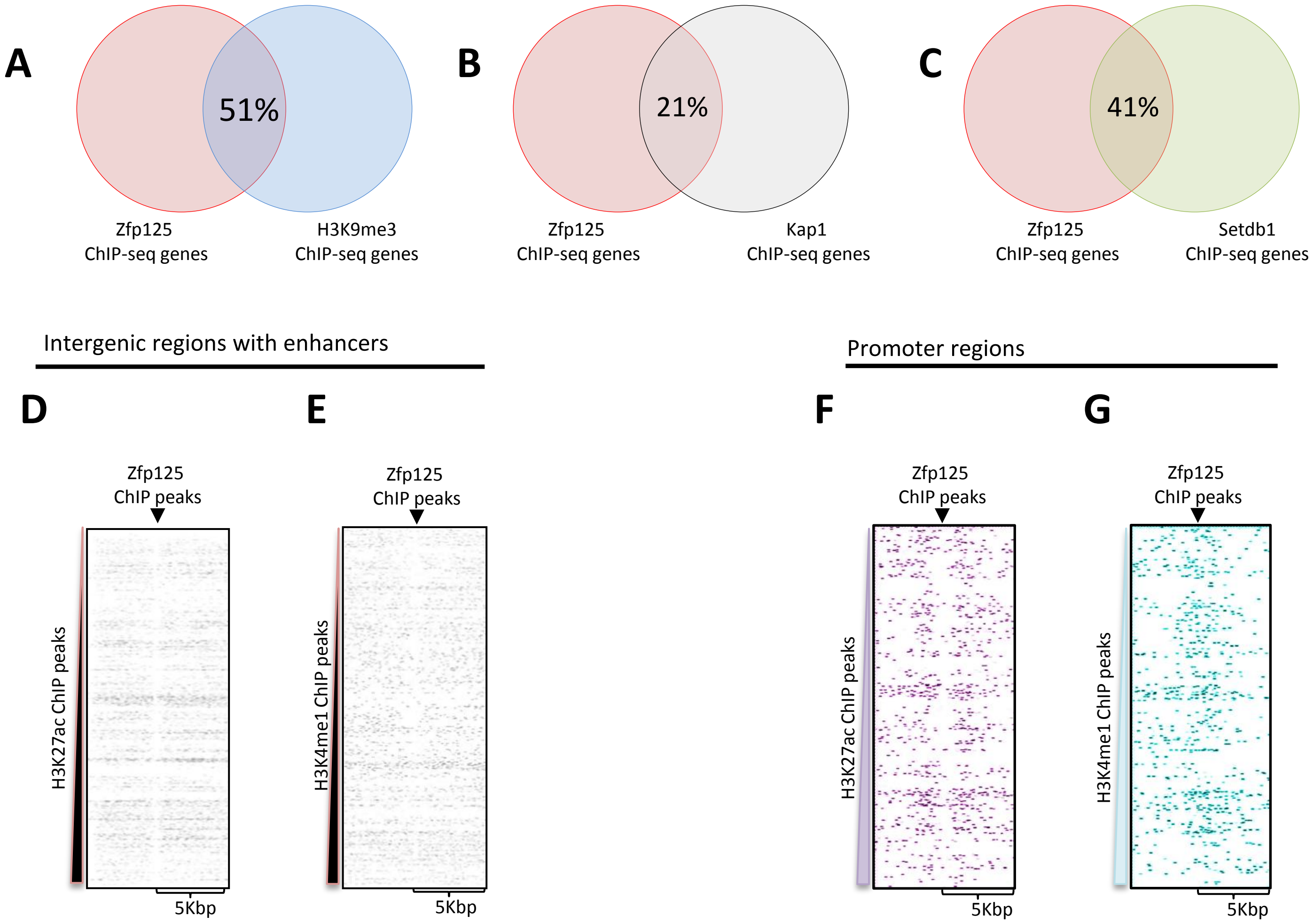
A
51%
Zfp125
ChIP-seq genes
H3K9me3
ChIP-seq genes
B
21%
Zfp125
ChIP-seq genes
Kap1
ChIP-seq genes
C
41%
Zfp125
ChIP-seq genes
Setdb1
ChIP-seq genes
Intergenic regions with enhancers
Promoter regions
D
E
F
G
Zfp125
ChIP peaks
H3K27ac ChIP peaks
H3K4me1 ChIP peaks
5Kbp

Figure S6

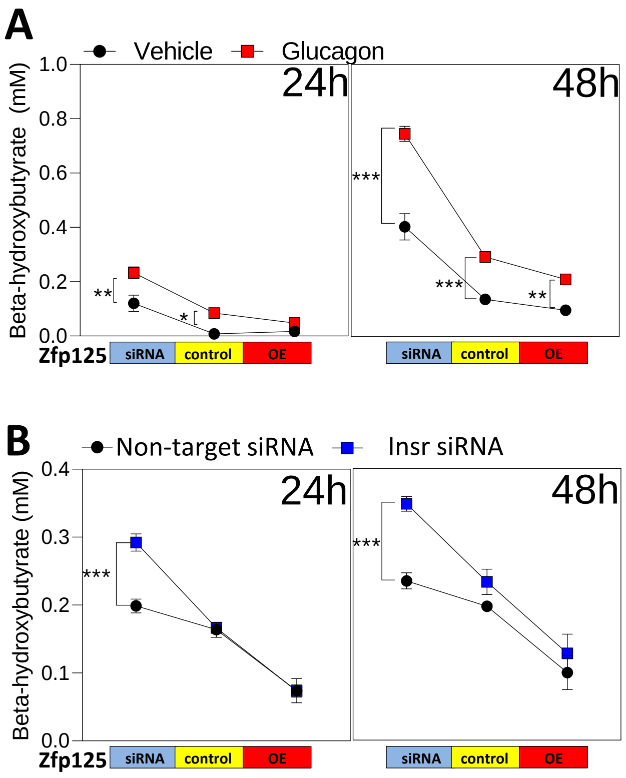
A
Vehicle
Glucagon
24h
48h
Beta-hydroxybutyrate (mM)
1.0
0.8
0.6
0.4
0.2
0.0
**
*
***
***
**
Zfp125
siRNA
control
OE
siRNA
control
OE
B
Non-target siRNA
Insr siRNA
24h
48h
Beta-hydroxybutyrate (mM)
0.4
0.3
0.2
0.1
0.0
***
***
Zfp125
siRNA
control
OE
siRNA
control
OE

Figure S7
